# *GBA1* mutations alter neuronal firing and structure, regulating VGLUT2 and CRYAB in dopamine neurons

**DOI:** 10.1101/2024.08.05.606574

**Authors:** Eva Rodríguez-Traver, Luz M. Suárez, Carlos Crespo, Irene González-Burgos, Rebeca Vecino, Juan C. Jurado-Coronel, María Galán, Marta González-González, Rosario Moratalla, Carlos Vicario

## Abstract

Mutations in the *GBA1* gene are major risk factors for Parkinsońs disease (PD), but their role in PD pathology is not fully understood. The impact of *GBA1* mutations was investigated in dopamine (DA) neurons obtained from induced pluripotent stem cells (iPSCs) derived from PD patients carrying the N370S or L444P *GBA1* mutation. DA neurons co-expressing TH and VGLUT2 were detected in the cultures, and their number and/or expression of *VGLUT2*/*SLC17A6* mRNA was markedly reduced in both N370S and L444P cultures compared to controls. A significant increase in the firing rate of N370S neurons was found, whereas evoked dopamine release was stronger from neurons carrying either mutation. Furthermore, mutant neurons accumulated abundant degenerative structures, and there was a significant accumulation of α-synuclein aggregates in N370S neurons. Notably, a significant upregulation of the chaperone *CRYAB/HSPB5/alpha-crystallin-B* was found early in DA neuron differentiation and in the substantia nigra of PD patients. Our findings indicate that N370S and L444P *GBA1* mutations impair midbrain DA neurons expressing VGLUT2, and provoke molecular, functional and structural changes, possibly involved in PD pathology.

## Introduction

The *GBA1* gene encodes the lysosomal enzyme β-glucocerebrosidase 1 (glucosylceramidase beta, GCase1), which catalyzes the hydrolysis of glucosylceramide (glucocerebroside or GlcCer) to ceramide and glucose, and that of glucosylsphingomyelin. Mutations in *GBA1* are among the strongest genetic risk factors for Parkinsońs disease (PD) (Baden et al, 2019; Blandini et al, 2019; Bloem et al, 2021; Cavallieri et al, 2023; Chatterjee & Krainc, 2023; Cilia et al, 2016; Gegg et al, 2022; Poewe et al, 2020; Sidransky & Lopez, 2012; Sidransky et al, 2009). Around 350 different mutations in *GBA1* have been reported but the heterozygous N370S and L444P are those most frequently found in PD patients. *GBA1* mutations have been estimated to increase the risk of PD 5- to 20-fold, while 5-25% of PD cases carry a mutation in this gene (Blandini et al., 2019; Garcia-Sanz et al, 2021; Poewe et al., 2020; Rodriguez-Traver et al, 2019a; Rodriguez-Traver et al, 2019b; Seto-Salvia et al, 2012; Sidransky & Lopez, 2012). Although *GBA1*-associated PD (*GBA1*-PD) has similar symptomatology to idiopathic PD, *GBA1* mutations favor an earlier onset of the disease, and a higher frequency of cognitive impairment and dementia (Baden et al., 2019; Blandini et al., 2019; Jiang et al, 2020; Mata et al, 2016; Obeso et al, 2022; Schapira et al, 2017; Seto-Salvia et al., 2012). Importantly, the brains of idiopathic PD patients without *GBA1* mutations have less GCase and weaker GCase activity, probably due to the deleterious effect of aggregated α-synuclein (α-syn), a major component of Lewy bodies, on the enzyme (Blandini et al., 2019; Fares et al, 2021; Mazzulli et al, 2011; Shahmoradian et al, 2019; Shults, 2006; Surmeier et al, 2017).

The molecular mechanisms that link the alterations to GCase1 protein structure and activity due to *GBA1* mutations with an increased risk of PD are still not fully understood. Several hypotheses have been proposed that associate GCase1 deficiency with α-syn aggregation. Firstly, a deficiency in GCase would lead to the accumulation of GlcCer in the lysosome, which could in turn perturb lipid homeostasis. This would stabilize α-syn oligomers, promoting the formation and accumulation of insoluble α-syn fibrils that would impair GCase trafficking from the endoplasmic reticulum (ER) to the Golgi apparatus (GA), further reducing enzyme activity in the lysosome (Garcia-Sanz et al., 2021; Garcia-Sanz et al, 2017; Mazzulli et al., 2011; Sidransky & Lopez, 2012; Yap et al, 2011; Zunke et al, 2018). Elsewhere, a direct interaction between GluCer and α-syn has been suggested that would lead to protein aggregation (Zunke et al., 2018). In fact, mutated GCase1 has been seen to co-localize with α-syn in lysosomes, Lewy bodies and neurites (Goker-Alpan et al, 2010). Moreover, it was recently proposed that α-syn degradation could be impaired by mutant GCase, leading to α-syn accumulation. Indeed, GCase misfolding in the ER leads to a loss of enzymatic activity that impedes α-syn degradation, which could drive the degeneration and death of DA neurons due to lysosomal-autophagic dysfunction (Horowitz et al, 2022). Along similar lines, recent evidence indicates that *GBA1* mutations impair α-syn degradation by inhibiting chaperone-mediated autophagy (CMA) (Kuo et al, 2022). Alternatively, reducing GCase activity may exacerbate pre-existing α-syn aggregation and hence, the pathological susceptibility of neurons (Henderson et al, 2020).

Thus, misfolded GCase alone, or in combination with α-syn aggregation, can generate dysfunctional phenotypes in the cells, inducing ER stress, activation of the unfolded protein response (UPR), lysosome and autophagic dysfunction, cholesterol accumulation, mitochondrial dysfunction, reactive oxygen species (ROS) production and defects in calcium homeostasis (Baden et al., 2019; Beccano-Kelly et al, 2023; Burbulla et al, 2017; Fernandes et al, 2016; Garcia-Sanz et al., 2017; Garcia-Sanz et al, 2018; Gegg et al., 2022; Horowitz et al., 2022; Lang et al, 2019; Mazzulli et al., 2011; Pradas & Martinez-Vicente, 2023; Schondorf et al, 2014; Schondorf et al, 2018; Stojkovska et al, 2021; Woodard et al, 2014; Zunke et al., 2018). All these findings illustrate the progress made in understanding the cellular and molecular mechanisms leading to cellular dysfunction and neuronal degeneration in *GBA1*-PD.

However, important questions remain to be addressed in order to gain a deeper understanding of the relationship between impaired GCase and PD etiopathology, which could shed light on potential treatments to slow or impede disease progression. In particular, it is fundamental to define the alterations triggered by *GBA1* mutations in DA neurons, those common to and those that distinguish each mutation, as well as to explore the extent to which these mutations affect both mature and developing neurons, or different DA neuron subtypes. Here we identified common and distinct molecular, functional and structural impairments in iPSC-DA neurons carrying N370S or L444P *GBA1* mutations. Importantly, some of these alterations appear in relatively immature DA neurons, such as reduced VGLUT2 and increased CRYAB expression, suggesting that they may potentially serve as molecular targets and biomarkers in *GBA1*-PD.

## Results

### The molecular phenotype of DA neurons derived from PD patients carrying mutations in *GBA1* and healthy controls

We initially assessed whether general neuronal and midbrain DA markers were expressed by cells during iPSC differentiation. In cultures of control cells and PD cells carrying N370S/wt or L444P/wt *GBA1* mutations (Fig. EV1), *PAX6* and *TUBB3* expression was detected identifying neural progenitors and immature neurons. Expression of genes encoding transcription factors (*LMX1A, LMX1B, FOXA2, and NURR1*), enzymes (*TH*), monoamine transporters (vesicular monoamine transporter2 - *VMAT2* - and dopamine transporter - *DAT*), and the G-protein-coupled inwardly-rectifying potassium channel 2 (*GIRK2*), which are molecular markers of midbrain DA neuronal progenitors and/or neurons (Andersson et al, 2006; Chu et al, 2002; Friling et al, 2009; Garritsen et al, 2023; Kamath et al, 2022; Kittappa et al, 2007; Lee et al, 2010; Mendez et al, 2005; Reyes et al, 2012; Rodriguez-Traver et al, 2016; Sanchez-Danes et al, 2012) was also detected. No significant differences in transcript relative levels were observed between PD and control cells (Fig. EV1A-B), although *TUBB3* and *DAT* expression was significantly weaker in N370S/wt cells than L444P/wt cells (*P<0.05; **P<0.01: Fig. EV1C). *GBA1* expression was similar in the three cell types.

After immunostaining cells with antibodies specific to TH, and the general neuronal markers β-III-tubulin (TUJ1) (Supplementary Fig. S1) and MAP2 (Fig. EV2), no significant differences were evident between the control (wt/wt) and PD cultures (N370S/wt or L444P/wt GBA1), with 47.1-56.0% TH^+^ neurons relative to the TuJ1^+^ neurons in the cultures at 30 DIV (Supplementary Fig. S1J), or 24.0-28.2% relative to MAP2^+^ neurons at 50 DIV (Fig. EV2J) and 40.4-49.0% at 90 DIV (Fig. EV2T). TH^+^ neurons began expressing the ventral dopaminergic marker FOXA2 early (47-53% FOXA2^+^/TH^+^ neurons at 21 DIV: Fig. EV3A-J), like the midbrain DA neuron marker GIRK2 (43.0-47.0% GIRK2^+^/TH^+^ neurons at 54 DIV: Fig. EV3K-T), with no significant differences found between the control and PD cultures.

### A subpopulation of DA neurons exhibits a glutamatergic phenotype: Effect of N370S and L444P *GBA1* mutations on these neurons

The presence of GABAergic and glutamatergic neurons in the cultures was also analysed, as they are found in the substantia nigra (Garritsen et al., 2023; Rodriguez & Gonzalez-Hernandez, 1999; Tritsch et al, 2012; Trudeau et al, 2014). The proportion of TH^+^ neurons that were also GABA^+^ was between 5.7% and 7.3%, with no significant differences between control and PD cultures (Supplementary Fig. S2A-J).

The vesicular glutamate transporter2 (VGLUT2) is expressed by nearly all midbrain DA neurons during embryonic development (Dumas & Wallen-Mackenzie, 2019; Eskenazi et al, 2021; Kouwenhoven et al, 2020; Steinkellner et al, 2018). Although in the adult substantia nigra its expression only persists in <20% of these neurons (Garritsen et al., 2023; Pereira Luppi et al, 2021; Steinkellner et al., 2018; Trudeau et al., 2014), VGLUT2 may be neuroprotective in PD (Buck et al, 2022; Hnasko et al, 2010; Kashani et al, 2007; Shen et al, 2018; Steinkellner et al, 2021). To determine if a subpopulation of hiPSC-derived DA neurons adopt a glutamatergic phenotype and whether this might be affected by the *GBA1* mutations, cultures were dual immunostained with antibodies against VGLUT2 and TH (Fig. 1A-I). The proportion of TH^+^ neurons co-expressing VGLUT2 was 6.6-fold lower in L444P/wt cultures (2.45% ±1.08) than in wt/wt cultures (16.12% ±6.03, P<0.01: Fig. 1J). There was also a clear trend towards fewer relative VGLUT2^+^/TH^+^ neurons in the N370S/wt cultures than in the wt/wt cells, although this was not statistically significant (Fig. 1J). Importantly, the percentage of DA neurons co-expressing VGLUT2 in the controls (16.12%) was similar to that in vivo (Steinkellner et al., 2018). The *SLC17A6/VGLUT2* mRNA expression by cells differentiating for 18 days was similar in N370S/wt and in L444P/wt cultures relative to the control cultures (Fig. 1K). However, the effect of each mutation on *VGLUT2* mRNA was reflected by the reduced *VGLUT2* expression in N370S/wt cultures relative to the L444P/wt cultures (**P<0.01: Fig. 1L).

**Figure 1.**
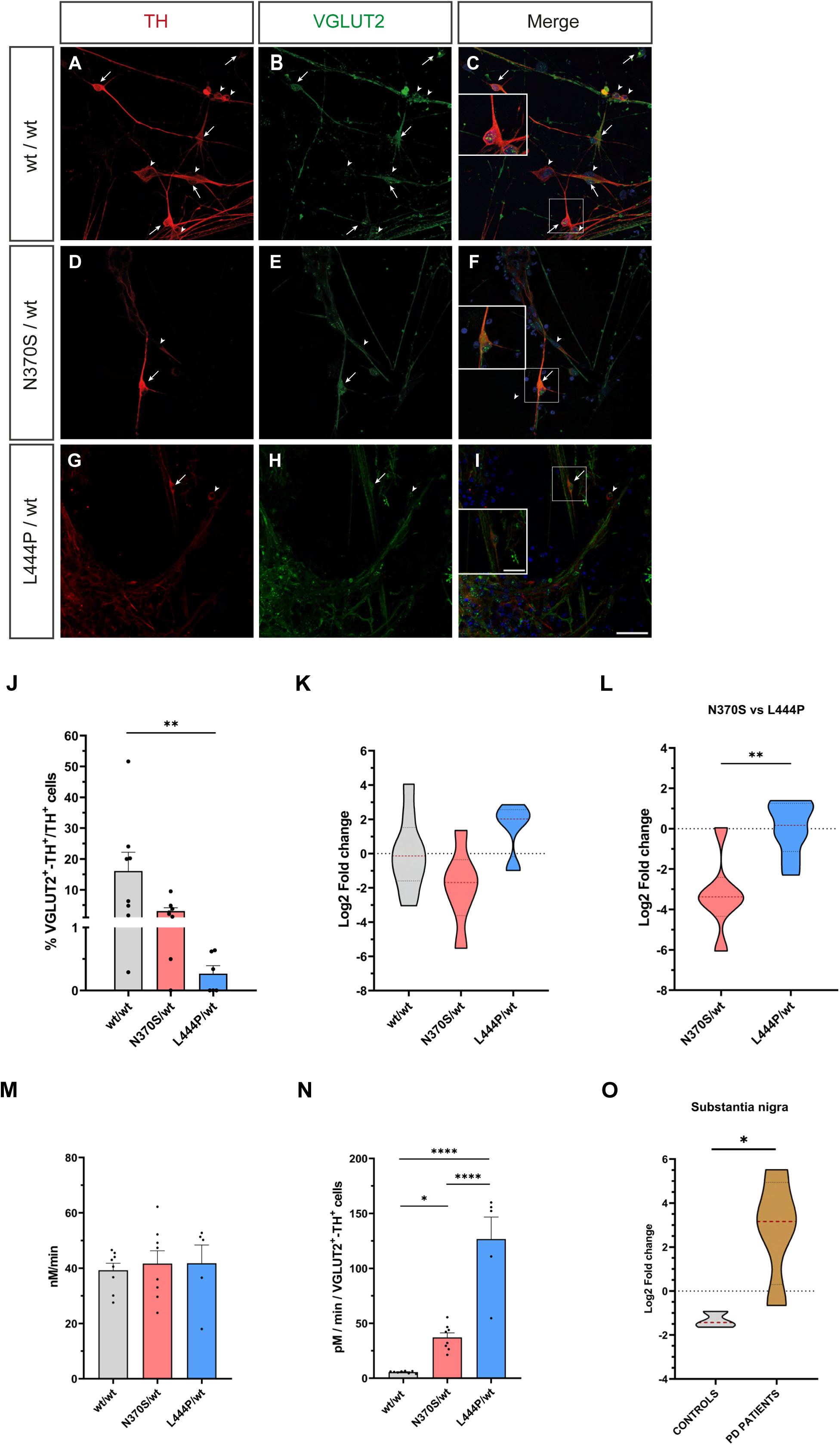
N370S and L444P *GBA1* mutations affect VGLUT2-DA neurons and *SLC17A6/VGLUT2* mRNA expression. (A-J) Neurons differentiating from human iPSCs were fixed at 50 DIV, and then immunostained with specific antibodies against TH and VGLUT2. The arrows indicate VGLUT2^+^/TH^+^ neurons and arrowheads point to TH^+^ neurons not expressing VGLUT2. (J) The percentage of TH^+^ neurons expressing VGLUT2 was strongly reduced in N370S/wt and L444P/wt cultures, although the reduction was only significant in L444P/wt relative to the control cultures (\*\**P*<0.01). (K, L) Analysis of *SLC17A6*/*VGLUT2* mRNA expression by RT-qPCR on 18 DIV cultures shows significantly weaker expression in N370S/wt than in L444P/wt cultures (\*\**P*<0.01). The data was analysed using an unpaired Student’s t test to compare the mean (±SEM values) of *n* = 4-6 cultures from 3 experiments performed in triplicate. (M) The level of glutamate (nM/min) in the culture medium was similar in all three conditions, although it was significant higher in N370S/wt and L444P/wt cultures relative to the number of VGLUT2^+^/TH^+^ neurons (\**P*<0.05; \*\*\*\**P*<0.0001: N). (O) *SLC17A6/VGLUT2* expression was significantly higher in the patient’s substantia nigra compared to that of the controls (*P<0.05). The data was analysed using an unpaired Student’s t test to compare the mean (±SEM values) of samples from 4 controls and 4 PD patients carrying no *GBA1* mutations, performed in triplicate. The data on the cell numbers and glutamate content are the mean (±S.E.M.) of *n* = 5-8 cultures per genotype from 2-3 experiments. The statistical analysis was carried out using a Kruskal-Wallis test followed by a Dunn test (cell counts) or one-way ANOVA followed by a Tukey test (glutamate levels). Scale bar: 40 µm; insets, 20 µm.

Since VGLUT2 mediates glutamate co-release and storage by DA neurons into synaptic vesicles (SVs) (Hnasko et al., 2010), we measured the glutamate in the culture medium. Although the total glutamate concentration was similar in all three conditions (Fig. 1M), when this concentration was expressed relative to the number of VGLUT2^+^-TH^+^ neurons (Fig. 1N), there were significant increases (7-23-fold) in the N370S/wt and L444P/wt cultures relative to the control cultures (*P<0.05; ****P<0.0001). Moreover, the glutamate concentration per VGLUT2^+^-TH^+^ neuron was significantly higher in L444P/wt than in N370S/wt cultures (****P<0.0001: Fig. 1N). Together, these findings suggest that the lower proportion of TH^+^ neurons expressing VGLUT2 and the increase in glutamate could enhance neuronal vulnerability in *GBA1*-associated PD.

RNA from the substantia nigra of PD patients and from healthy control subjects was then analysed, and we detected that *VGLUT2* mRNA expression was significantly higher in the patient’s substantia nigra compared to that of the controls (*P<0.05: Fig. 1O).

### The impact of *GBA1* mutations on dopamine release

Dopamine was detected in the cultures under both basal conditions (Fig. 2A) and after KCl-induced depolarization (evoked: Fig. 2B). There was no significant difference in dopamine content between *GBA1*-PD and control cultures. The dopamine released per minute in wt/wt, N370S/wt and L444P/wt cultures was 50, 104 and 146 times greater in evoked than in basal conditions (Fig. 2C). These increases were significant in N370S/wt (*P<0.05) and L444P/wt cultures (****P<0.001), yet not in the control cultures. Moreover, there was a significant increase in the released dopamine under evoked conditions in the L444P/wt cultures relative to the control cultures (^####^P<0.0001: Fig. 2C). Hence, iPSC-derived DA neurons carrying *GBA1* mutations respond more strongly to depolarization than control neurons.

**Figure 2.**
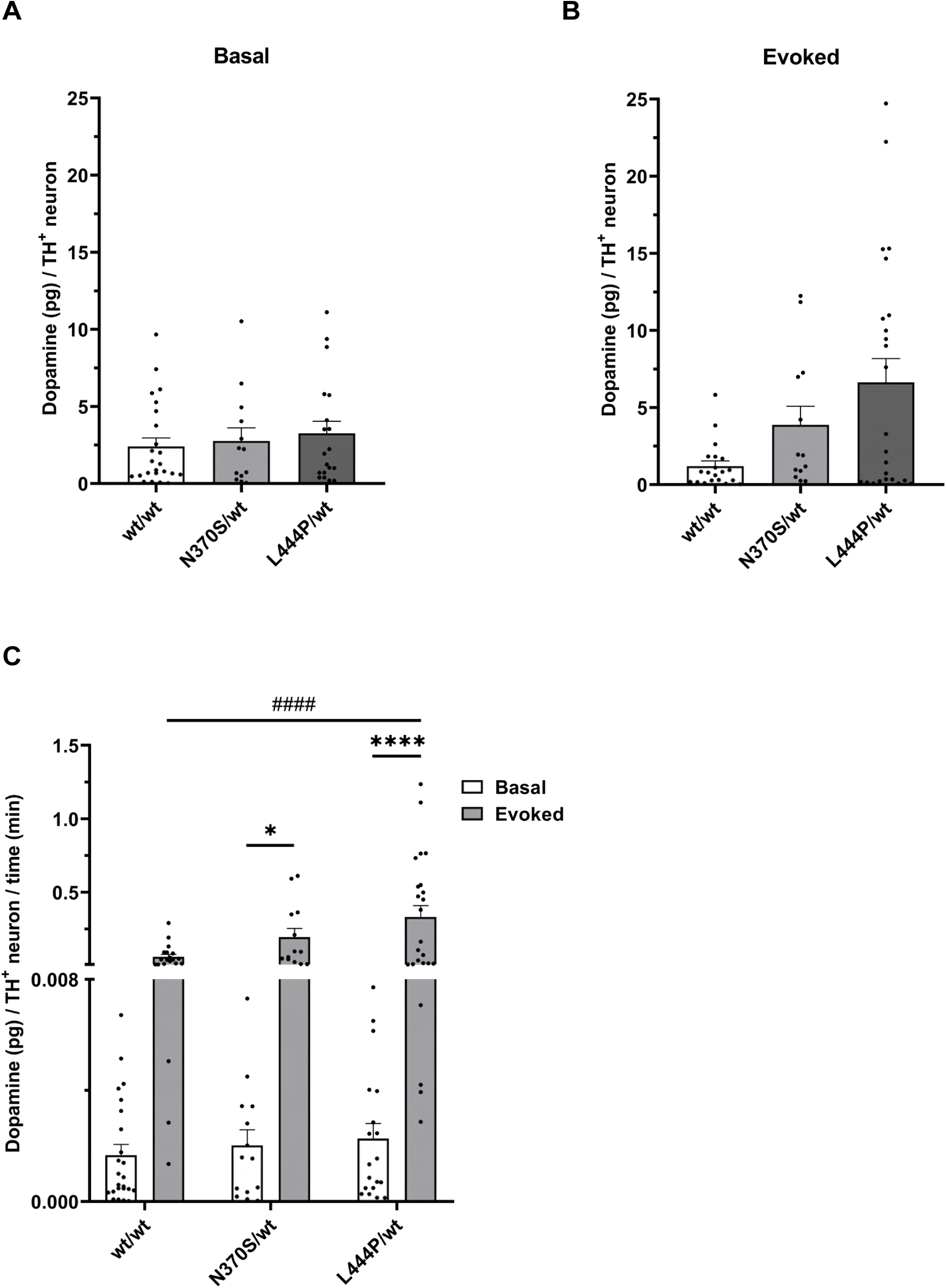
The N370S and L444P *GBA1* mutations alter dopamine release in response to KCl. Basal (A) and KCl-evoked (B) dopamine release by iPSC-derived DA neurons into the culture medium was measured by ELISA after 47, 51 and 59 DIV, combining the results in the analysis. The data are expressed as dopamine (pg)/TH^+^ neuron (A, B) or as dopamine (pg)/TH^+^ neuron per minute (C), and presented as the mean (±S.E.M.) of *n* = 13-24 cultures per genotype from 3 experiments. The KCl-evoked dopamine release was significantly stronger than that of the basal dopamine release in N370S/wt and L444P/wt cultures (\**P*<0.05; \*\*\*\**P*<0.0001), but this was not significantly different in the wt/wt cultures. The evoked dopamine release was significantly higher in the L444P/wt than in wt/wt cultures (^####^*P*<0.0001). Statistical analysis was performed using one-way ANOVA with a Tukey post-hoc test.

### The effect of *GBA1* mutations on the electrical responses of DA neurons

It was important to demonstrate that iPSC-derived DA neurons were electrically active and to assess whether the *GBA1* mutations modified the electrophysiological properties of these neurons. DA neurons were identified by ICC using antibodies against TH and MAP2 after they were recorded, filled with Alexa-Fluor 488 and fixed (Supplementary Fig. S3A-F and Fig. S4A-L). *GBA1* mutations did not significantly alter the passive properties of DA neurons (Supplementary Fig. S3G-J). In terms of active properties (Fig. EV4A), a first action potential (AP) was evoked using depolarizing current pulses in 75% of

DA neurons in the wt/wt cultures, as opposed to 100% of N370/wt and L444P/wt DA neurons, although this increase was not significant. Furthermore, more mutant neurons responded with a second AP than wt/wt neurons, although there were no differences between the control and GBA1 mutant neurons in terms of the threshold, peak amplitude and half amplitude duration of APs (Fig. EV4B-E). After injecting hyperpolarizing and depolarizing current pulses, the I-V curves had a similar pattern in all three conditions (Fig. EV4F). Intrinsic repetitive firing of APs was detected, indicating the presence of neurons with the autonomous pacemaking properties typical of substantia nigra DA neurons (Fig. EV4G) (Guzman et al, 2009; Li et al, 2022; Pu et al, 2023) However, the effect of *GBA1* mutations on this repetitive firing was not addressed here.

Notably, the firing rate of N370S/wt DA neurons was significantly higher than that of the control neurons (*P<0.05; **P<0.01; ***P<0.001: Fig. 3A), meaning that these neurons generated more APs per second for the same injected current. The L444P/wt neurons appeared to have a slightly but not significantly higher firing rate than control neurons. Hence, iPSC-derived DA neurons acquire functional features and N370S/wt DA neurons adopt a hyperexcitable electrical phenotype. Finally, spontaneous excitatory postsynaptic currents (sEPSCs) were analysed in voltage-clamp mode (Fig. 3B), with more mutant neurons developing sEPSCs than control (wt/wt) neurons, although the differences were not significant (Fig. 3C). The amplitude of the sEPSCs was similar in all three conditions (Fig. 3D), yet the frequency of sEPSCs was significantly lower in L444P/wt neurons than in controls (**P<0.01: Fig. 3E).

**Figure 3.**
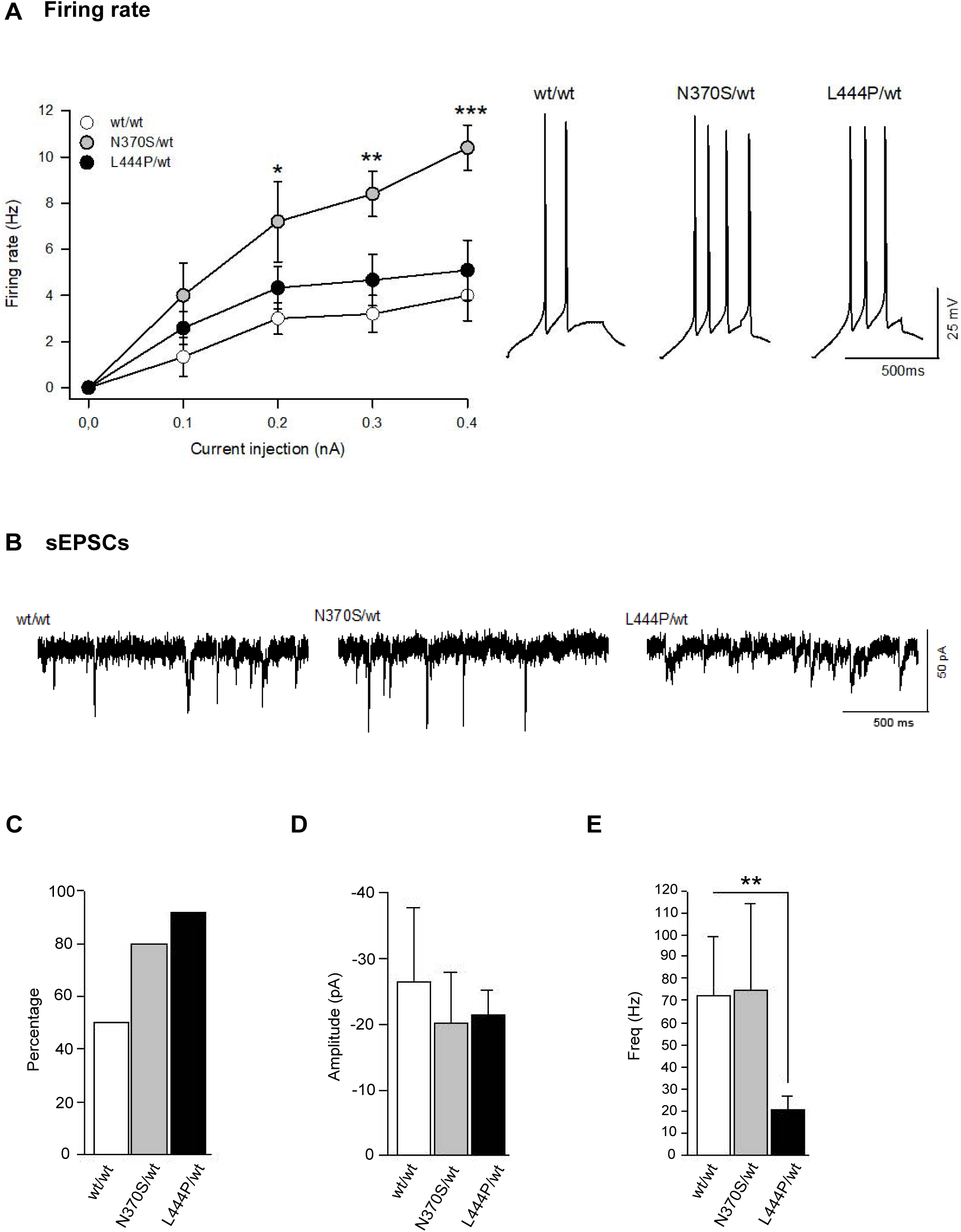
The N370S *GBA1* mutation increases the electrical firing rate of DA neurons. The human iPSC-derived neurons from control subjects and PD patients with N370S/wt or L444P/wt mutation in *GBA1* were maintained in culture for 90-93 days to perform electrophysiological recordings. The neurons recorded were injected with Alexa 488 and their dopaminergic phenotype was confirmed by immunostaining for TH (see supplementary Fig. S3A-F and S4A-L). (A) The firing rate increased significantly in N370S/wt neurons relative to wt/wt neurons. (B) The traces illustrating spontaneous excitatory postsynaptic currents (sEPScs) are shown. The *GBA1* mutations increased, albeit not significantly, the proportion of neurons that developed sEPSCs (C) and produced no changes in their amplitude (D). However, significantly fewer sEPSCs were detected in L444P/wt neurons relatve to wt/wt neurons. The results are the mean (±S.E.M.) of *n* = 6-12 neurons per genotype from 2 experiments: \**P*<0.05; \*\**P*<0.01; \*\*\**P*<0.001 (two-way ANOVA followed by a Bonferronís test).

Our morphological analysis of the recorded TH^+^ cells showed that L444P/wt DA neurons had significantly more intersections in the first 60 µm from the cell body than control and N370S/wt neurons (*P<0.05; **P<0.01; ***P<0.001: supplementary Fig. S4A-P). There were no other appreciable changes in morphological parameters (Supplementary Fig. S5A-C).

### The N370S *GBA1* mutation favours α-synuclein accumulation in DA neurons

α-synuclein is enriched in the presynaptic compartment, and it influences vesicle dynamics, trafficking and neurotransmitter release (Calabresi et al, 2023; Zalon et al, 2024). However, due to its unstable conformation it may undergo misfolding under cellular stress and aggregate in the PD brain, leading to the formation of Lewy bodies (Alam et al, 2019; Goker-Alpan et al., 2010; Shahmoradian et al., 2019; Zalon et al., 2024). Thus, we assessed whether α-syn accumulates in the cell body and dendrites of iPSC-derived DA neurons (Fig. 4). Neurons carrying N370S/wt mutation had significantly more aggregates in both cell bodies (**P<0.01: Fig. 4A-M) and dendrites (****P<0.0001: Fig. 4N-Z) relative to wt/wt neurons. However, the number of aggregates in L444P/wt neurons was similar to those in control neurons and significantly lower than in N370S/wt neurons, both in the soma (***P<0.001: Fig. 4M) and dendrites (****P<0.0001: Fig. 4Z). Furthermore, the N370S/wt culture medium contained more α-syn per TH^+^ neuron than that of wt/wt and L444P/wt cultures, although this difference was not significant (Fig. 4Z’). Hence the two *GBA1* mutations do not have the same effect on the mechanisms driving α-syn aggregation and release by iPSC-derived DA neurons.

**Figure 4.**
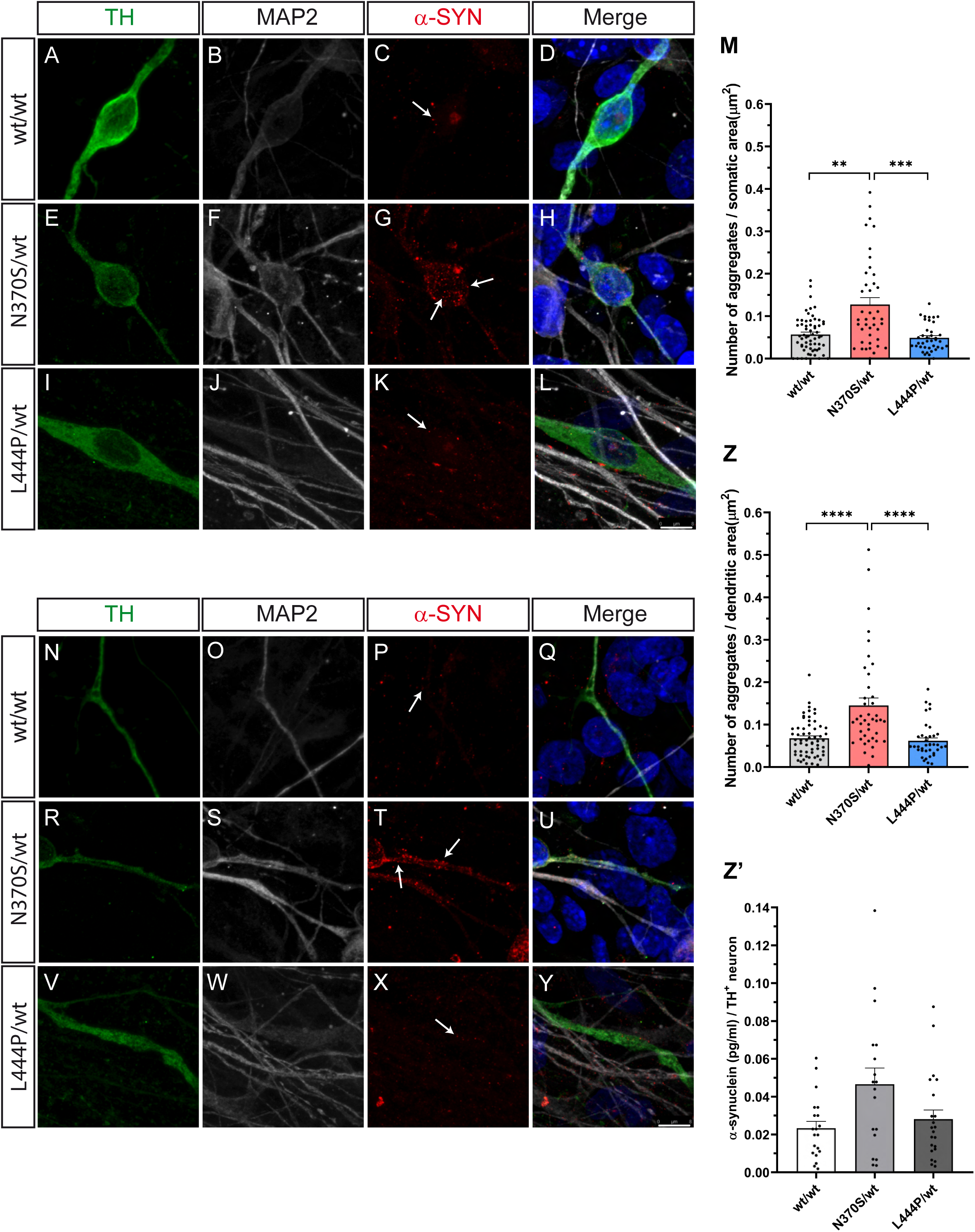
The N370S *GBA1* mutation augments the α-synuclein aggregates in cell bodies and dendrites of DA neurons. (A-L, N-Y) Human iPSC-derived neurons from control subjects and PD patients carrying N370S/wt or L444P/wt mutations in *GBA1* were maintained in culture for 95 or 103 days and then immunostained for TH, MAP2 and α-synuclein. The aggregates in the soma (M) and dendrites (Z) of TH^+^ neurons were semi-automatically counted using ImageJ after tracing the neuron and setting the thresholds. Arrows point to examples of aggregates. The results are the mean (±S.E.M.) of *n* = >40 neurons per genotype from 3 experiments and they are expressed as the number of aggregates per soma or per dendritic area. There were significantly more α-synuclein aggregates in the cell bodies and dendrites of N370S/wt *GBA1*-DA neurons than in wt/wt and L444P/wt neurons: \*\**P*<0.01; \*\*\**P*<0.001; \*\*\*\**P*<0.0001 (Kruskal-Wallis test followed by Dunńs multiple comparisons test). Scale bar: 8 µm. There was a trend towards more α-synuclein in the culture medium in N370S/wt *GBA1*-cultures, albeit not significant (Z’).

### The N370S and L444P *GBA1* mutations alter the ultrastructure of iPSC-derived neurons

We then decided to use TEM to determine whether the presence of *GBA1* mutations alters the ultrastructure of iPSC-neurons cultured for 90-104 days. However, this analysis could not be specifically performed on TH^+^ neurons for technical reasons. In contrast to control neurons (Fig. 5A-B), structures resembling degenerative bodies were frequently detected in neurons derived from patients carrying the N370S (Fig. 5C-D) or L444P (Fig. 5E-F) *GBA1* mutations. These structures were full of vacuoles resembling autophagosomes and dysfunctional lysosomes. In fact, autophagosomes, autophagosome-like structures, and autophagosome-like vacuoles, were clearly distinguished in the cell bodies and processes of neurons carrying *GBA1* mutations (Supplementary Fig. S6B-D), which were barely detected in control neurons (Supplementary Fig. S6A). Multilamellar bodies (MLBs) were also abundant in mutant neurons (Fig. 6) and the membrane of the MLB’s were more disorganized in N370S (Fig. 6A-C) than in L444P neurons (Fig. 6D-H).

**Figure 5.**
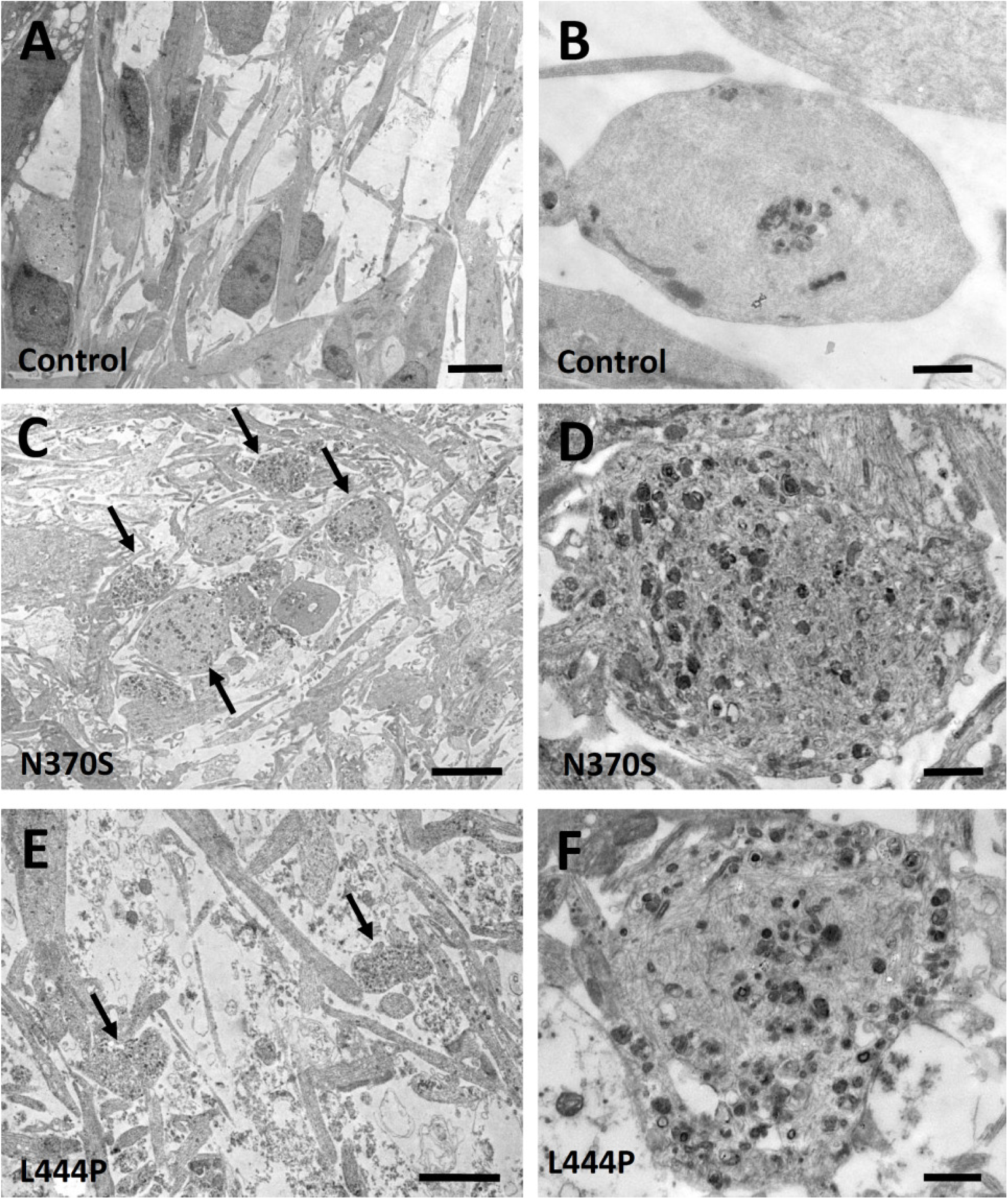
Abundant degenerative bodies form in N370S and L444P *GBA1* mutant neurons. (A, C, and E) Low-magnification micrographs from 104 DIV cultures of iPSC-derived neurons from control healthy subjects (A) and from patients carrying the N370S/wt (C) or the L444P/wt (E) *GBA1* mutations. Note that mutant neurons contain large structures that resemble degenerative bodies (arrows in C and E). (B, D and F) Micrographs of degenerative bodies at higher magnification. (B) This cell derived from a control subject contain a few autophagosome-like vacuoles. The large degenerative body-like structures found in patient-derived neurons carrying the N370S/wt (D) or the L444P/wt (F) mutation are full of vacuoles resembling autophagosomes and dysfunctional lysosomes. Scale Bars: A, C, E, 5 µm; B, D, F, 1 µm.

**Figure 6.**
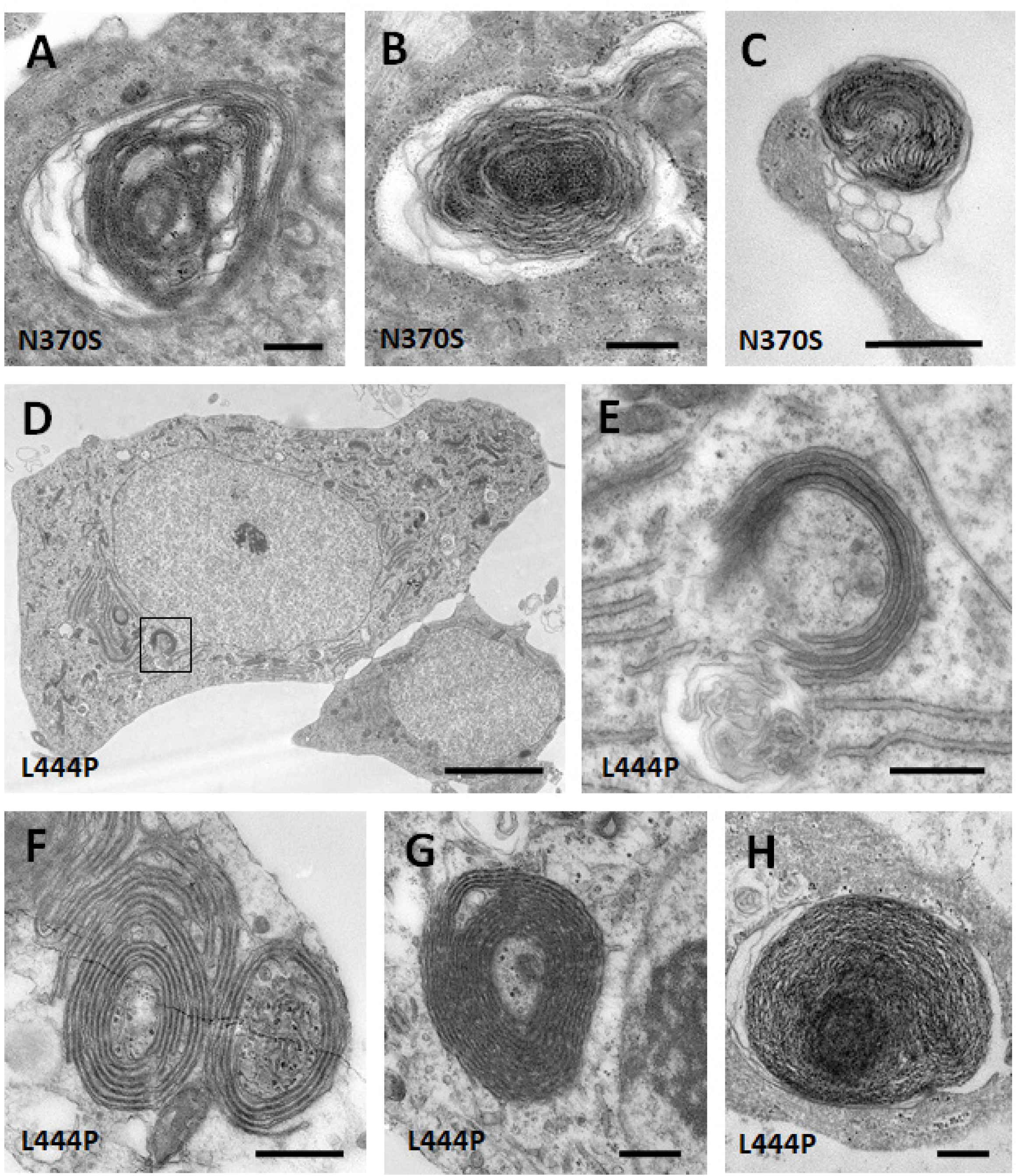
The N370S and L444P *GBA1* mutations promote multilamellar body formation in neurons. (A-C) Multilamellar bodies (MLBs) in cultured iPSC-derived neurons from patients carrying the N370S/wt *GBA1* mutation. The membranes that integrate these structures are clearly disorganized. (D) Low magnification micrograph showing a cultured iPSC-derived neuron from a patient carrying the L4444P/wt *GBA1* mutation. The boxed area is shown at higher magnification in (E). This cell contains an ‘atypical’ structure of membrane cisternae arranged in a semi-circular manner. Note the electron-dense material located among the cisternae. (F-H) Multilamellar structures in cultured iPSC-derived neurons from patients carrying the L444P/wt *GBA1* mutation. The membrane cisternae that integrate the structures in F and G are perfectly distinguishable. Note that these multilamellar structures enclose cytoplasmic remnants with heterogeneous material. The membranes that integrate the structure shown in H are disorganized. Scale Bars: A-C, E-H, 500 nm; D, 5 µm.

Since mutant GCases are retained in the GA (Fernandes et al., 2016; Garcia-Sanz et al., 2018; Mazzulli et al., 2011; Sidransky & Lopez, 2012; Zalon et al., 2024), this organelle was also analysed (Fig. 7). In fact, whereas GA dictyosomes had a normal appearance in control iPSC-derived neurons (Fig. 7A), large numbers of abnormal and vacuolated GA dictyosomes were evident in neurons derived from patients carrying the N370S (Fig. 7B-C) or L444P *GBA1* mutation (Fig. 7D-F). Indeed, while the percentage of vacuolated dictyosomes in control cells was 10.25%, this number increased to 42.85% and to 82.60% in cells carrying the N370S or the L444P mutation, respectively (Fig. 7G).

**Figure 7.**
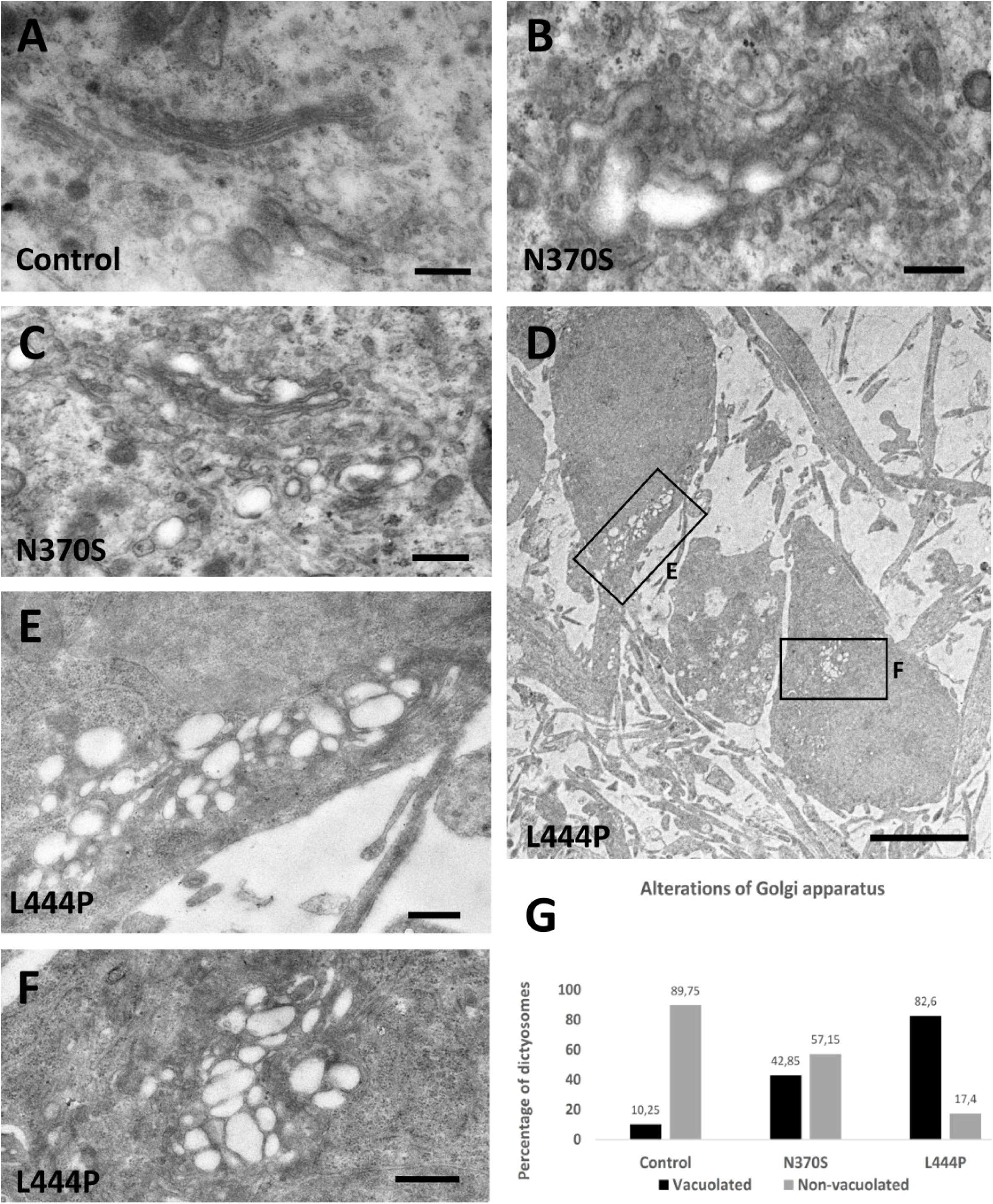
The N370S and L444P *GBA1* mutations alter the Golgi apparatus of iPSC-derived neurons. (A) Golgi apparatus (GA) dictyosomes have a normal appearance in iPSC-derived neurons from control healthy subjects. (B-F) A large number of GA dictyosomes with an abnormal and vacuolated appearance was found in iPSC-derived neurons from patients carrying the N370S/wt (B, C) or L444P/wt *GBA1* mutation (D-F). The dictyosomes in the boxed areas in D are shown at higher magnification in E and F. Note that these dictyosomes are completely vacuolated. (G) Graph depicting the proportion of vacuolated and non-vacuolated dictyosomes in iPSC-derived neurons from controls and from patients carrying the N370S/wt or L444P/wt *GBA1* mutation. Scale Bars: A, 4 µm; B, C, 500 nm; D, E, F, 250 nm.

In summary, both N370S and L444P *GBA1*-neurons accumulate abundant degenerative bodies, MLBs, vacuolated GA dictyosomes, autophagosomes and dysfunctional lysosomes that could affect neuronal function and vulnerability.

### Expression of *CRYAB* in DA neurons and in the substantia nigra of PD patients

Small chaperones fulfil several roles, such as acting as molecular sensors of cellular stress or preventing protein misfolding (Hayashi & Carver, 2020) and mutations in the *GBA1* gene cause alterations in GCase1 folding (Kuo et al., 2022; Patnaik et al, 2012; Sidransky & Lopez, 2012). We previously found an upregulation of *CRYAB* (or *HSPB5/alpha-crystallinB*) in neural stem cell cultures (Nieto-Estevez et al, 2013) and hypothesized that this chaperone might be expressed by iPSC-derived DA neurons, especially in those derived from the patients. Immature neurons (30 DIV) from wt/wt individuals and from patients carrying the N370S/wt and L444P/wt *GBA1* mutations expressed CRYAB protein in both their soma and processes (Fig. 8A-I). CRYAB was detected in 25.12% wt/wt TH^+^ neurons, whereas nearly twice as many N370S/wt TH^+^ neurons expressed CRYAB (47.65% cells; P<0.0001: Fig. 8J) and 39.19% of the TH^+^ L444P/wt cells were CRYAB^+^ (a non-significant increase of 56% relative to the control neurons). As further insight into the molecular mechanism involved, RT-qPCR analysis revealed significantly stronger *CRYAB* mRNA expression in N370S/wt cultures than in wt/wt (**P<0.01: Fig. 8K) or L444P/wt cultures (*P<0.05: Fig. 8L), indicating that the N370S/wt mutation alters *CRYAB* transcription at early phases of DA neuron differentiation. Next, RNA from the substantia nigra of PD patients and from healthy controls was obtained, and *CRYAB* mRNA expression was seen to be significantly enhanced in the patients relative to the controls (*P<0.05: Fig. 8M), whereas no differences DAT, TH or *GBA1* mRNA expression were observed (Supplementary Fig. S7A). Furthermore, the expression of the five transcripts studied was similar in the hippocampus of controls and PD patients (Supplementary Fig. 7B).

**Figure 8.**
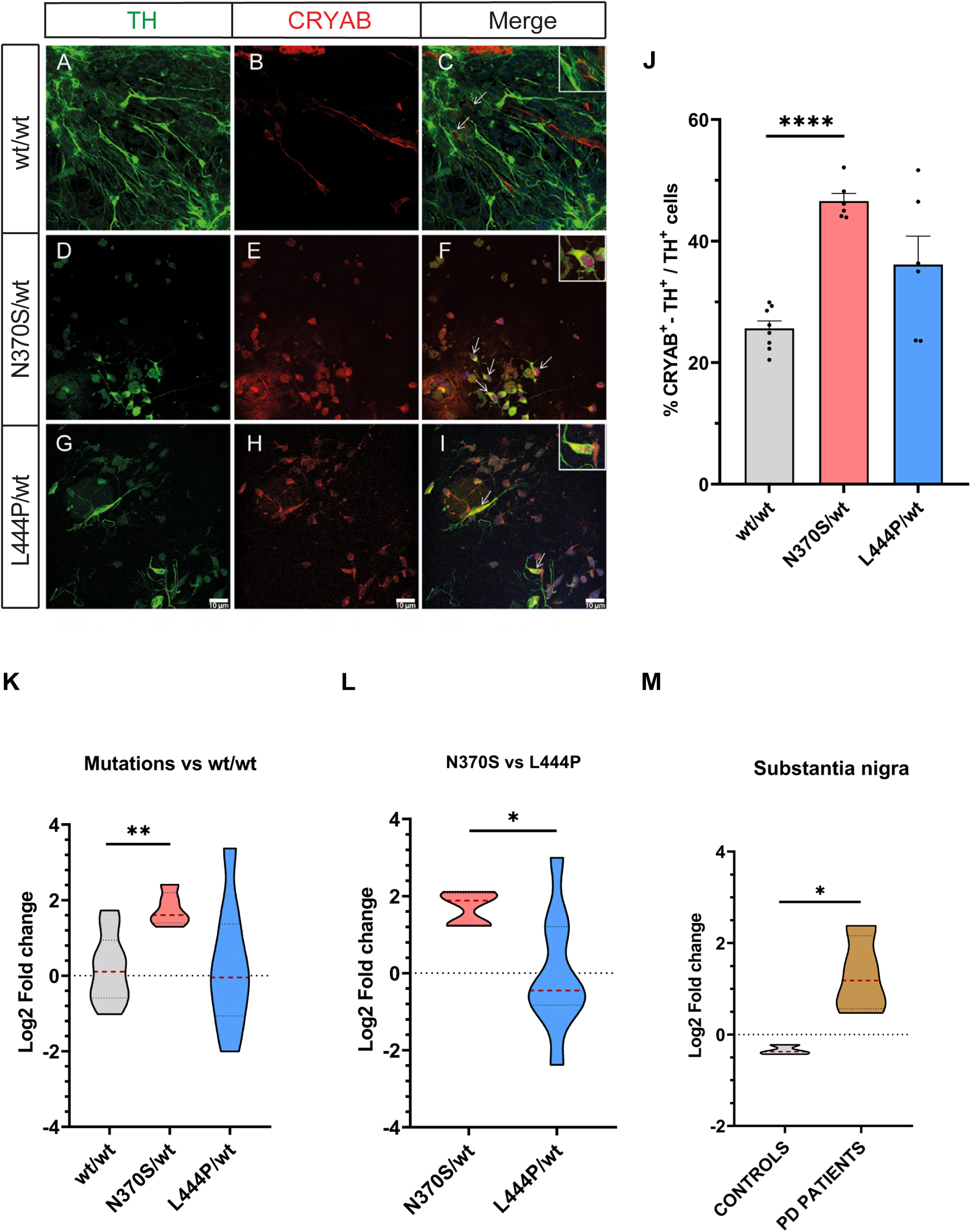
The N370S *GBA1* mutation increases CRYAB expression in DA neurons. (A-I) Human iPSC-derived cells from control subjects and PD patients carrying N370S/wt or L444P/wt mutations in *GBA1* were maintained in culture for 30 days and then immunostained for TH and CRYAB. (J) There was a marked increase in the DA neurons expressing CRYAB in the N370S/wt cultures relative to the controls (****P<0.0001), while the increase in L444P/wt cultures was not statistically significant. The arrows indicate CRYAB^+^/TH^+^ neurons and the results are the mean (±S.E.M.) of *n* = 6-8 cultures per genotype from 2-3 experiments. The statistical analysis was carried out using one-way ANOVA followed by Games-Howell’s post-hoc test. Scale bar: 10 µm. (K, L) Analysis of *CRYAB* (*HSPB5/alpha-crystallin-B*) mRNA expression by RT-qPCR in samples of 18 DIV cultures. A significant increase in *CRYAB* expression was found in N370S/wt cultures relative to the controls (\*\**P*<0.01) and to the L444P/wt cultures (\**P*<0.05). The statistical analysis was carried out using an unpaired Student’s t test with Welch correction to compare the mean (±S.E.M.) values of *n* = 4-6 cultures from 3 experiments. (M) *CRYAB* expression was significantly higher in the patient’s substantia nigra compared to that of the controls (\**P*<0.05). The data was analysed using an unpaired Student’s t test to compare the mean (±SEM values) of samples from 4 controls and 4 PD patients carrying no *GBA1* mutations, performed in triplicate.

## Discussion

The cellular and molecular mechanisms leading to cell dysfunction and neuronal degeneration in *GBA1*-PD are gradually becoming better understood (Baden et al., 2019; Brooker et al, 2024; Chatterjee & Krainc, 2023; Fernandes et al., 2016; Garcia-Sanz et al., 2021; Garcia-Sanz et al., 2018; Mazzulli et al, 2016; Pradas & Martinez-Vicente, 2023), although important issues are still to be resolved. Here we show that hiPSC-neurons carrying the N370S or the L444P *GBA1* mutation in heterozygosis show both relatively common (neuron structure, dopamine release) and distinct (electrical firing rate, α-syn aggregation, VGLUT2 and CRYAB expression) dysfunctional and degenerative features. While both mutations negatively affect neuronal ultrastructure, some features like MLB structure and the number of vacuolated GA dictyosomes are partially mutation-dependent. Importantly, CRYAB expression is enhanced in N370S immature neurons, as also observed in the substantia nigra of PD patients, evidence that CRYAB may possibly serve as an early biomarker and/or molecular target for PD. Furthermore, we report novel findings suggesting that N370S and L444P *GBA1* mutations could trigger DA neuron dysfunction and degeneration by dampening VGLUT2 mRNA and/or protein expression.

### VGLUT2 expression in DA neurons is altered by *GBA1* mutations

Analysis by RT-qPCR and immunostaining indicates that the N370S mutation is associated with significantly weaker *VGLUT2* mRNA expression, whereas there are fewer VGLUT2^+^-TH^+^ neurons in L444P cultures. The fact that the two mutations have partially different effects on *VGLUT2* expression, and on the proportion of DA neurons expressing this glutamate transporter, suggests that they may trigger some distinct actions at the molecular level. These could include direct effects on mRNA transcription, post-translational regulation and/or the activation of protein degradation (see below). To the best of our knowledge this is the first time VGLUT2 has been seen to be potential molecular targets of *GBA1* mutations during the development of human mDA neurons.

In mice, the large majority of embryonic mDA neurons express VGLUT2 (Dumas & Wallen-Mackenzie, 2019; Eskenazi et al., 2021; Kouwenhoven et al., 2020). Although similar studies have not been performed in human embryos, our data suggest that *GBA1* mutations may affect the glutamatergic and dopaminergic tone of the extracellular milieu of developing DA neurons. Thus, reduced VGLUT2 expression could enhance the vulnerability of DA neurons as VGLUT2 has been reported to be neuroprotective in PD, probably because this transporter promotes glutamate and dopamine storage in SVs (Buck et al., 2022; Hnasko et al., 2010; Kashani et al., 2007; Steinkellner et al., 2021; Trudeau et al., 2014). Our finding that *VGLUT2* mRNA is more strongly expressed in the substantia nigra of PD patients than in controls may reflect a role for this transporter in the resilience to neurodegeneration. However, no direct relationship could be established between *VGLUT2* and *GBA1 in vivo* as the post-mortem tissue analysed was from patients who do not carry *GBA1* mutations, although our data on cultured iPSC-derived DA neurons indicates that *GBA1* mutations affect VGLUT2 expression.

### The N370S and L444P *GBA1* mutations produce a hyperexcitable phenotype in DA neurons

Here we demonstrate that the iPSC-derived DA neurons (from both controls and PD patients) express features of mature and functional neurons, which include the extracellular release of dopamine under KCl stimulation, AP firing, and spontaneous postsynaptic currents. The fact that dopamine release in response to KCl is stronger in cultures from PD patients than from controls indicates that mutant DA neurons are more sensitive to depolarization. Because VGLUT2 promotes dopamine storage in SVs, stronger release may partially be due to the lower proportion of VGLUT2^+^/TH^+^ neurons in mutant *GBA1* cultures. Impaired dopamine release is a feature of PD (Cramb et al, 2023) and our findings suggest that this impairment can be visualized in culture through the increase in dopamine levels in response to KCl. Because dopamine can be converted into toxic oxidized dopamine (Song et al, 2023; Zhang et al, 2019), the hyperexcitable state of mutant DA neurons that drives stronger dopamine release could have deleterious consequences for these neurons. By contrast, we do not detect reduced basal dopamine levels in our cultures of DA neurons carrying *GBA1* mutations, consistent with earlier data (Schondorf et al., 2014). However, lower basal dopamine content was reported in iPSC-derived DA neurons from a N370S-GBA1 individual (Woodard et al., 2014) and in N370S-GBA1 brain organoids (Rosety et al, 2023). Among the considerations that must be borne in mind when interpreting these results is that the dopamine concentration was expressed as fmol/µl or ng/ml in earlier studies (Rosety et al., 2023; Woodard et al., 2014), whereas here it was expressed as pg/TH^+^ neuron because the amount of dopamine released is a function of the number of TH^+^ neurons in the culture.

The most striking and novel electrophysiological finding of our study is the hyperexcitable phenotype of N370S-*GBA1* neurons, with no changes in the activation threshold, suggesting that N370S neurons may have suffered changes in membrane ion channels. By contrast, multielectrode recording highlighted a deficiency in the spontaneous excitability of immature neurons derived from a PD patient carrying the N370S mutation. However, the firing rate of these N370S neurons increased with time in culture (Woodard et al., 2014) In midbrain organoids derived from N370S-*GBA1* patients, largely composed of neural progenitors and immature neurons, microelectrode arrays found reduced firing rates at 15 DIV (Rosety et al., 2023). The different protocols to obtain DA neurons and organoids, and the distinct differentiation/maturation stages at which recordings were made could partially account for the differences in these results. Here, mature N370S DA neurons (90-93 DIV) were patch-clamp recorded, whereas previous studies were carried out on N370S immature neurons (30 and 52 DIV) (Woodard et al., 2014) and on N370S organoids (15 DIV) (Rosety et al., 2023).

Importantly, higher firing frequencies have been reported in DA neurons from the substantia nigra *in vivo* after proteasome inhibition (Subramaniam et al, 2014). Since α-syn aggregates can affect the ubiquitin-proteasome system (Calabresi et al., 2023; Lindersson et al, 2004; Onal et al, 2024; Stojkovska et al., 2021; Zalon et al., 2024), α-syn accumulation in N370S-neurons could be involved in promoting hyperexcitability. In this sense, increased firing frequencies in DNAJC6 mutant midbrain-like organoids, a model of early-onset PD, coincide with α-syn aggregation (Wulansari et al, 2021). Here, electrical hyperexcitability was only evident in N370S-*GBA1* neurons, suggesting a correlation between increased α-syn aggregation and altered electrical firing, as reported in other neuronal systems (Calabresi et al., 2023; Tomagra et al, 2023; Tozzi et al, 2021; Wulansari et al., 2021).

### The N370S and L444P *GBA1* mutations disrupt neuronal ultrastructure, and they produce different effects on α-synuclein and CRYAB

Our EM study shows that neurons carrying *GBA1* mutations develop clear signs of degeneration, accumulating degenerative bodies, MLBs and vacuolated Golgi dictyosomes, with these features being partially mutation-specific. Although some of these features were previously observed in N370S fibroblasts, such as MLBs (Garcia-Sanz et al., 2017; Garcia-Sanz et al., 2018), this is the first time they are reported in human neurons derived from *GBA1*-PD patients. Therefore, our culture system allows clear features of neurodegeneration to be detected in iPSC-derived neurons, in contrast to the idea that this process is little evident in these human neurons.

Moreover, the accumulation of α-syn aggregates in TH^+^/MAP2^+^ neurons carrying the N370S mutation supports the feasibility of using this cell model to study neurodegeneration in PD. This accumulation is consistent with a previous study where such aggregates were also detected by ICC (Woodard et al., 2014), although ultrastructural alterations were not reported. L444P neurons did not accumulate significant α-syn or release this protein, suggesting that the ultrastructural defects in these neurons occur in the absence of α-syn aggregates, or alternatively, that the aberrant α-syn structures that might have formed in L444P neurons are rapidly degraded. By contrast, α-syn accumulation was previously detected in Western blots of L444P cell extracts (Schondorf et al., 2014) and the overexpression of this mutation in neurons was sufficient to induce lysosomal dysfunction (Mazzulli et al., 2016; Zalon et al., 2024). The α-syn accumulation in N370S-mutant neurons could be a consequence of impaired CMA, which is more deficient in iPSC-derived DA neurons carrying N370S than L444P mutation (Kuo et al., 2022; Onal et al., 2024). Together, our findings suggest that the aggregation and the degradation of aberrant proteins could be affected distinctly by the two *GBA1* mutations in the context of PD. Notably, we found a higher proportion of DA neurons expressing CRYAB and *CRYAB* mRNA upregulation in cultures from N370S PD patients. This protein may protect against the aggregation, and toxicity of amyloidogenic proteins (Liu et al, 2015), and it has been shown to participate in the inhibition of α-syn accumulation (Bruinsma et al, 2011; Cox et al, 2014; Masilamoni et al, 2006; Scheidt et al, 2021). However, CRYAB has not been previously described as an ER-associated and/or UPR-associated chaperone in cellular models of neurodegeneration (Fernandes et al., 2016; Onal et al., 2024; Zalon et al., 2024). The upregulation of *CRYAB* mRNA in the substantia nigra but not the hippocampal cortex of PD patients suggests CRYAB could be specifically involved in PD. Furthermore, the higher abundance of *CRYAB* mRNA and of CRYAB^+^/TH^+^ neurons at early stages of iPSC differentiation, well before α-syn aggregation in N370S neurons, suggest that CRYAB upregulation may act as a stress sensor in early stages of the neurodegenerative process.

**In conclusion**, our study uncovers common and distinct effects of the N370S and L444P *GBA1* mutations in multiple processes in neurons, for the first time identifying VGLUT2 and CRYAB as potential molecular targets and/or biomarkers of early stages of GBA1-associated PD.

## Methods

Please, see the Appendix (Supplementary material) for a full description of the Methods.

### Differentiation of human iPSCs (hiPSCs) into DA neurons

The vector-free hiPSCs used in this study were generated and characterized previously in our laboratory (Rodriguez-Traver et al, 2020; Rodriguez-Traver et al., 2019a; Rodriguez-Traver et al., 2019b). To differentiate the hiPSCs into DA neurons, we designed a protocol based on previous publications (Doi et al, 2014; Kirkeby et al, 2012; Kriks et al, 2011). First, the hiPSC colonies grown on mouse embryonic fibroblasts (MEFs) were dissociated into a single cell suspension by incubating them with accutase. For differentiation, the pellet of hiPSC was resuspended in MEF-conditioned hiPSC medium supplemented with FGF-2 (6 ng/mL) and Y-27632 (10 μM). The cell suspensions were plated on vitronectin-coated culture dishes at a density of 50,000-80,000 cells/cm^2^, and the medium was changed every other day. Once the cultures reached a cell confluence of 90-100%, neural induction commenced by changing the medium to hiPSC medium supplemented with Noggin (250 ng/mL, day 0), containing A83-01 (5 μM) or SB431542 (10 µM) in fewer experiments. The medium was changed on day 1 of the culture to hiPSC medium supplemented with Noggin, A83, Sonic Hedgehog C25II (SHH, 100 ng/mL), Purmorphamine (2 μM) and FGF-8 (100 ng/mL). On day 3, CHIR99021 (CHIR, 3 μM) was also added along with the factors added on day 1. From day 5 to day 9, the hiPSC medium was incrementally shifted to DMEMF12N2 medium (25%, 50%, 75%) every 2 days, and on day 7 and 9 the medium was only supplemented with Noggin and CHIR. On day 11, the medium was changed to Neurobasal/B27/GlutaMax supplemented with CHIR and with BDNF (20 ng/mL), ascorbic acid (0.2 mM), GDNF (20 ng/mL), dibutyril cAMP (0.5 mM) and TGFβ3 (1 ng/mL: BAGCT compounds). From day 13, CHIR was no longer added, BAGCT was included and the medium was changed every other day until the cells were dissociated with accutase and collected on days 17-20 (depending on the confluence of the cultures). The cells recovered were then plated on coated coverslips in the wells of 24 multi-well plates, or on 48, 12 or 6 multi-well plates at 300,000-400,000 cells/cm^2^. For long-term maturation studies, cells were plated on a layer of mitotically inactivated primary astrocytes at 150,000-200,000 cells/cm^2^. Depending on the studies carried out, the cells were maintained in culture from 18 to 104 days. Control, N370S-*GBA1* and L444P-*GBA1* hiPSCs were always differentiated in parallel, and similarly, all the assays and analyses were performed in parallel.

### Real-Time Quantitative Reverse Transcription PCR (RT-qPCR)

The NucleoSpin RNA extraction kit (Macherey-Nagel) was used to purify the RNA from the cells, according to the manufacturer’s instructions. Complementary DNA (cDNA) was obtained by reverse transcription of 250-500 ng of the initial RNA using the SuperScript III enzyme. To determine the relative expression of different genes the RT-qPCR analysis was performed in triplicate using Power SYBR Green or Power SYBR Green Fast and the primer pairs were designed with either Lasergene software or purchased directly (Supplementary Table S1). Changes in gene expression were compared between the N370S/wt and L444P/wt cultures, and also with the expression in the control (wt/wt) cultures, using the comparative C_T_ method with the equation 2^-ΔΔC^ (Nieto-Estevez et al., 2013; Pfaffl, 2001; Schmittgen & Livak, 2008). Values were obtained as the fold change on a log2 scale, and then represented using violin plots after conversion to a linear scale. Statistical analysis was performed using an unpaired Student’s t test, applying Welch’s correction when the F-test indicated that the variances of both groups differed significantly.

### Immunocytochemistry (ICC)

Cells were fixed for 25 minutes with 4% paraformaldehyde (PFA) and then incubated for 1 hour at room temperature (RmT) in permeabilization/blocking buffer (0.1-0.4% Triton X-100/Normal goat or donkey serum/PBS), prior to probing them 20-22 hours with primary antibodies against: CRYAB (1:250: Abcam Cat# ab76467, RRID:AB_1523120); FOXA2 (1:100: Abcam Cat# ab108422, RRID:AB_11157157); GABA (1:2000: Sigma Cat# A2051, RRID:AB_2314459); GIRK2 (1:200: Abcam Cat# ab65096, RRID:AB_1139732); MAP2 (1:250: Sigma-Aldrich Cat# M1406, RRID:AB_477171 and 1:1000: Synaptic Systems Cat# 188004, RRID:AB_2138181); α-synuclein LB509 (1:800: Abcam Cat# ab27766, RRID:AB_727020); TH (1:200: Millipore Cat# MAB318, RRID:AB_2201528 and 1:200: Millipore Cat# AB152, RRID:AB_390204); β-III-tubulin (TUJ1, 1:1000: Covance Cat# MMS-435P, RRID:AB_2313773 and TUJ1, 1:300: Abcam Cat# ab18207, RRID:AB_444319); VGLUT2 (1:500: Alomone Cat# AGC-036, RRID:AB_2340950). Antibody binding was then detected with the appropriate secondary antibodies for 1 h at RmT.

For quantification, confocal images from 5 or 10 random microscope fields were taken per culture using a 63x objective or a 40x objective with a 1.5 zoom. Cells in the entire Z-stack were counted and the co-localization of specific markers was analysed in each individual Z-plane using ImageJ software (NIH, Bethesda, MD). The results were expressed as the mean ± S.E.M. of 3-33 cultures per genotype from 3-7 differentiation experiments. The specific n values for each ICC assay are given in the figure legends and the statistical analysis was carried out using one-way ANOVA followed by a Bonferrani or Tukey post-hoc test, or Kruskal-Wallis test followed by a Dunn post-hoc test.

### Analysis of α-synuclein aggregates in DA neurons

Cells fixed on DIV 95 or 103 were probed for 20-22 h at 4 °C with antibodies against TH, MAP2 and α-syn, and then with the appropriate secondary antibodies. Confocal images (each containing a clearly defined TH^+^ neuron) were obtained with a 63x objective using a 3.5 zoom, and three images were taken from each neuron: one from the soma and two from two dendrites. The number of α-syn aggregates was semi-automatically counted using ImageJ.

First, a threshold for detecting TH^+^ staining was set to detect the neuron of interest which was then manually traced and its somatic and dendritic area measured. Next, a threshold for α-syn^+^ staining was applied to quantify the number of aggregates. This allowed for an automatic detection of aggregates above a specific threshold. The aggregates were determined blind to the genotype, and expressed as the number of aggregates per area. Statistical analysis was performed using the non-parametric Kruskal-Wallis test followed by Dunńs multiple comparisons test.

### Dopamine release

To measure the basal dopamine, the culture medium was replaced with fresh medium for 24 hours before it was collected. Subsequently, fresh medium with KCl (56 mM) was added to the cells and after 20 min it was collected to measure the dopamine release evoked, while the cells were fixed with 4% PFA for TH ICC to determine the total number of TH^+^ cells per well. Dopamine was extracted and measured using the Dopamine Research Elisa™ assay kit (Labor Diagnostika Nord GmbH & Co. KG). Calculations were made following a non-linear regression for curve fitting using GraphPad Prism 5.0. The dopamine data was expressed as picograms (pg) of dopamine per TH^+^ neuron, or as pg of dopamine per TH^+^ neuron and per minute, as the mean (±S.E.M.) of 13-24 cultures per genotype and from 5 experiments. Statistical analysis was performed using one-way ANOVA with a Tukey post-hoc test.

### Quantification of glutamate in the culture medium

Basal glutamate levels in the culture medium of iPSC-differentiating neurons (47 DIV) were determined using an enzymatic and colorimetric assay (Glutamate Assay Kit, Sigma Aldrich) after protein precipitation with 1M perchloric acid followed by neutralization with 2M KOH and centrifugation. Glutamate was determined by measuring the absorbance at 450 nm in a spectrophotometer (FLUOstar OPTIMA; BMG Labtech). The glutamate data was expressed as glutamate (nM)/min or as glutamate (pM/min)/VGLUT2^+^ - TH^+^ neurons, as the mean (±S.E.M.) of 5-8 cultures per genotype and from 2 experiments. Statistical analysis was performed using one-way ANOVA with a Tukey post-hoc test.

### Quantification of α-synuclein in the culture medium

The hiPSC derived neurons were maintained in culture for 47 or 90 DIV, when the medium was collected to determine the α-syn levels using the SimpleStep ELISA kits (Abcam). No α-syn was detected in the α-syn knockout cells (according to the manufacturer), reflecting the specificity of this assay. Calculations followed a near-linear regression for fitting using GraphPad Prism 5.0 software and fixed cells were immunostained with an antibody against TH. The data were expressed as picograms (pg/ml) of α-syn per TH^+^ neuron and are the mean (±S.E.M.) of 16-19 cultures per genotype from 2 experiments. Statistical analysis was performed using one-way ANOVA with a Tukey post-hoc test.

### Electrophysiology

Electrophysiological experiments were performed on hiPSC-differentiated neurons (90-93 DIV) cultured on mouse astrocytes growing on coverslips, which were transferred to a submersion-type recording chamber mounted on a Leica microscope. The chamber was continuously perfused (0.5 ml/min) with carbogen (95% O_2,_ 5% CO_2_)-bubbled artificial cerebrospinal fluid (aCSF) at 36.5 °C. The aCSF medium was adjusted to 310 mOsm with sucrose and it contained (in mM): 130 NaCl, 4 KCl, 2 CaCl_2_, 1 MgCl_2_, 10 HEPES, and 10 glucose [pH 7.4]. Whole-cell patch-clamp pipettes had an initial resistance of 5– 10 MΩ. The internal recording solution was adjusted to an osmolarity of 290 mOsm with sucrose and it contained (in mM): 130 Cs-gluconate, 10 NaCl, 2 MgCl_2_, 0.2 EGTA, 1 NaATP, and 10 HEPES adjusted to pH 7.2. To visualize the cells, Alexa Fluor-488 hydrazide sodium salt (125 µM) was added to the internal solution.

Patch-clamp recordings were obtained in the whole-cell configuration using Axoclamp 2-B (Molecular Devices) and digitized at 3 KHz using 1440A Digidata (Molecular Devices). The amplifier bridge circuit was adjusted to compensate for serial resistance and monitored throughout the experiment. Passive properties (resting membrane potential - RMP, membrane capacitance - Cm, and membrane resistance-Rm) were analysed using the membrane test function integrated in the pClamp10 software. Neuronal excitability was assessed in current clamp mode with a 500 ms somatic current injection and spontaneous synaptic activity was recorded holding the cells at −70 mV (voltage clamp). The data were stored and analysed on a digital system (pClamp v10.0, Molecular Devices). Results are presented as the mean (±S.E.M.) of 6-12 DA neurons (90-93 DIV) from two experiments. The data was analysed using one- or two-way ANOVA with a Dunn or Bonferroni post-test, respectively, or using the Chi-squared test.

### Morphological analysis of the cells recorded

Neurons were injected with Alexa Fluor-488 while electrophysiological recordings were carried out and subsequently, they were fixed with 4% PFA and immunostained with antibodies against TH and MAP2. The samples were scanned using an AF 6000LX Leica microscope with a 20x objective to identify the Alexa Fluor 488 labelled cells. TH^+^ neurons were traced using ImageJ Sholl analysis software using the Bonfire program in MATLAB (Kutzing et al, 2010). For statistical analysis, a two-way ANOVA with a Bonferronís test was performed. The average length of the processes, their total length and the number of processes was also quantified and analysed using a one-way ANOVA with a Tukeýs post-test. The results are expressed as the mean (±S.E.M.) of 8-9 neurons per genotype from 3 experiments.

### Electron microscopy

The hiPSC-differentiated neurons (90-104 DIV) cultured on a mouse astrocyte monolayer on coverslips were examined by transmission electron microscopy (TEM). The cells were fixed in 4% PFA and 2% glutaraldehyde in 0.12M Phosphate Buffer (PB, pH 7.4), and then treated for 45 minutes at RmT with 1% OsO_4_ (Electron Microscopy Sciences-EMS) and 7% glucose in PB, after which they were stained with 1% uranyl acetate (EMS) in maleate buffer (pH 4.5). The cells were then dehydrated in increasing concentrations of ethanol (50°, 70°, 96° and 100°), and washed twice for 10 minutes in 100° ethanol and CuSO_4_ to complete dehydration. After clearing in propylene oxide (Fluka AG) and embedding in Durcupan, the Durcupan was polymerized for 24 h at 60 °C. The next day, the coverslips were placed on top of pre-made resin columns and the resin was polymerized at at 60 °C for 24 h. Ultrathin sections (75–80 nm) were obtained and then studied under transmission TEM.

### Analysis of human samples

#### RNA isolation from human post-mortem tissue

Total RNA was isolated from post-mortem tissue (10-20 mg) from the substantia nigra and hippocampal cortex of PD patients or healthy subjects. None of these patients carried *GBA1* mutations and in fact, despite contacting many Brain Banks, none could provide us with tissue from *GBA1*-PD patients. The tissue was mechanically disrupted and homogenized 4-5 times with a Potter-Elvehjem tissue homogenizer in buffer RA1 (NucleoSpin® RNA kit) with β-mercaptoethanol. The samples were then passed 6 times through a 0.9 mm syringe needle, and passed through a filter to clear the lysate and reduce viscosity. The extraction protocol for the kit was then followed, assessing the concentration and quality of the RNA obtained using NanoDrop One.

#### Summary of the statistical analysis

After first determining whether the data followed a normal distribution using the Shapiro-Wilk test, parametric (unpaired t-test, t-test with Welch correction for different variances, one-way and two-way ANOVA) or non-parametric tests (Mann-Whitney and Kruskal-Wallis tests) were used for the statistical analysis. The ROUT method was applied to score data outliers and P<0.05 was considered significant in all the analysis performed to discard the null hypothesis. GraphPad Prism 5.0 and 8.0 was used for all the statistical analyses.

## Supporting information

Complete Methods and Supplemental figures

## Acknowledgements

We thank the following individuals from the Instituto Cajal-CSIC (Madrid, Spain): Dr Eva Díaz-Guerra and Dr Alicia Hernández-Vivanco for their help with the experiments; Paula Vicario del Río, Silvia Calero, Pablo Fernández, and Lucía Vicario del Río for their help with the image analysis and formatting the references; and Mª José Román for her technical support. We also thank Dr Mark Sefton (BiomedRed SL, Madrid, Spain) for English editing advice and the “Unidad de Imagen Científica y Microscopía” (Instituto Cajal-CSIC) for helping us with the analysis of α-syn aggregates. We appreciate the contribution of the responsible and coordinator personnel of the CIEN tissue bank (BT-CIEN, Fundación CIEN, Madrid, Spain) and CeGeN-ISCIII (Santiago de Compostela, Spain) for providing us with post-mortem tissue and performing the genotype analysis, respectively.

## Author contributions

E.R.-T.: Designed and performed cell culture experiments, cellular and molecular assays, data analysis and interpretation, manuscript revision. L.M.S.: Performed electrophysiology, data analysis and interpretation, manuscript revision. C.C.: Performed electron microscopy, data analysis and interpretation, manuscript revision. I.G.-B.: Performed cellular and molecular assays, data analysis and interpretation, manuscript revision. R.V.: Performed molecular assays, data analysis and interpretation. J.C.J.-C.: Performed cellular assays, data analysis and interpretation. M.G.: Performed cellular assays, data analysis and interpretation. M.G.-G.: Performed molecular assays, data analysis and interpretation. R.M.: Conceptualization, experimental design, data analysis and interpretation, manuscript revision, financial and administrative support. C.V.: Conceptualization, experimental design, cell culture, supervision, data analysis and interpretation, manuscript writing, revision and editing, financial and administrative support, final approval of manuscript.

## Ethical approval

The study was approved by the Commission of Guarantees for the Donation and Utilization of Human Cells and Tissues, Instituto de Salud Carlos III (ISCIII), Spain (approval numbers: 3212701, 4083311, 4103331 and 5224261). Informed consent was obtained from all the patients.

## Data availability statement

The data that support the findings of this study are available on reasonable request from the corresponding author.

## Supplementary information is available in the Appendix

### The Paper Explained Problem

Although the cause of Parkinsońs disease is largely unknown, mutations in the *GBA1* gene are major risk factors for this disease. However, the available treatments do not stop the progression of Parkinsońs disease.

## Results

Our study on dopamine neurons obtained from human stem cells shows that *GBA1* mutations alter the structure, excitability, and protein expression in these neurons.

### Impact

We identify the proteins VGLUT2 and CRYAB as potential molecular targets and/or biomarkers of early stages of *GBA1*-Parkinsońs disease, which could shed light on potential treatments to slow or impede disease progression.

## FIGURE LEGENDS (Expanded View figures)

**Figure EV1.**
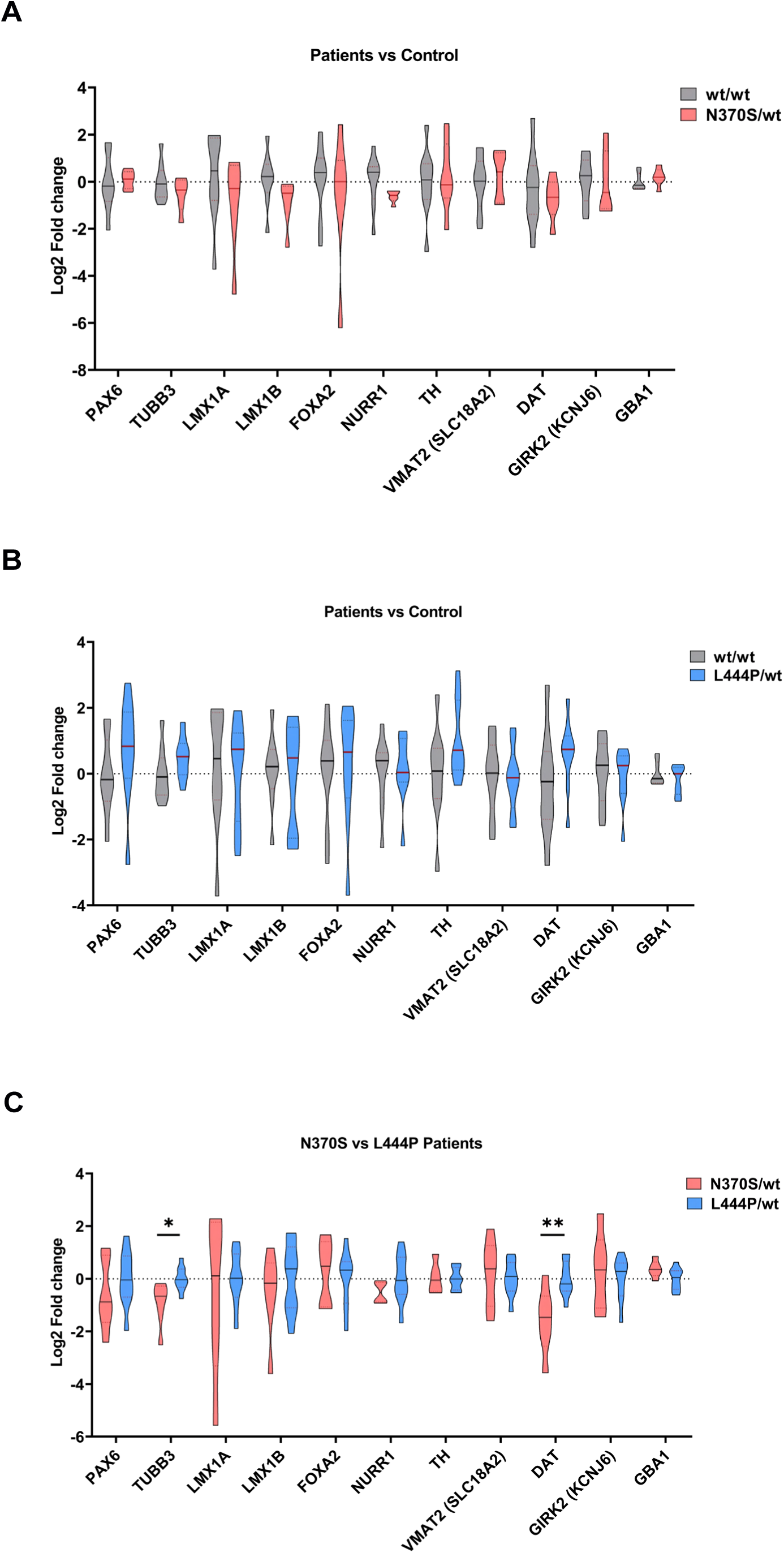
Gene expression in differentiating cultures of iPSCs derived from *GBA1*-PD patients and healthy subjects. (A-C) Total RNA was extracted from the cells after 18 days of differentiation, reverse transcribed to cDNA and analysed by RT-qPCR using the oligonucleotide primers listed in Supplementary Table S1. Transcript expression was largely similar in the three conditions, although a significantly lower expression of *TUBB3* and *DAT* was found in N370S/wt relative to the L444P/wt cultures (\**P*<0.05; \*\**P*<0.01: C). Statistical analysis was carried out using an unpaired Student’s t test to compare the mean (±SEM) values of *n* = 6-9 cultures per genotype from 3 experiments performed in triplicate.

**Figure EV2.**
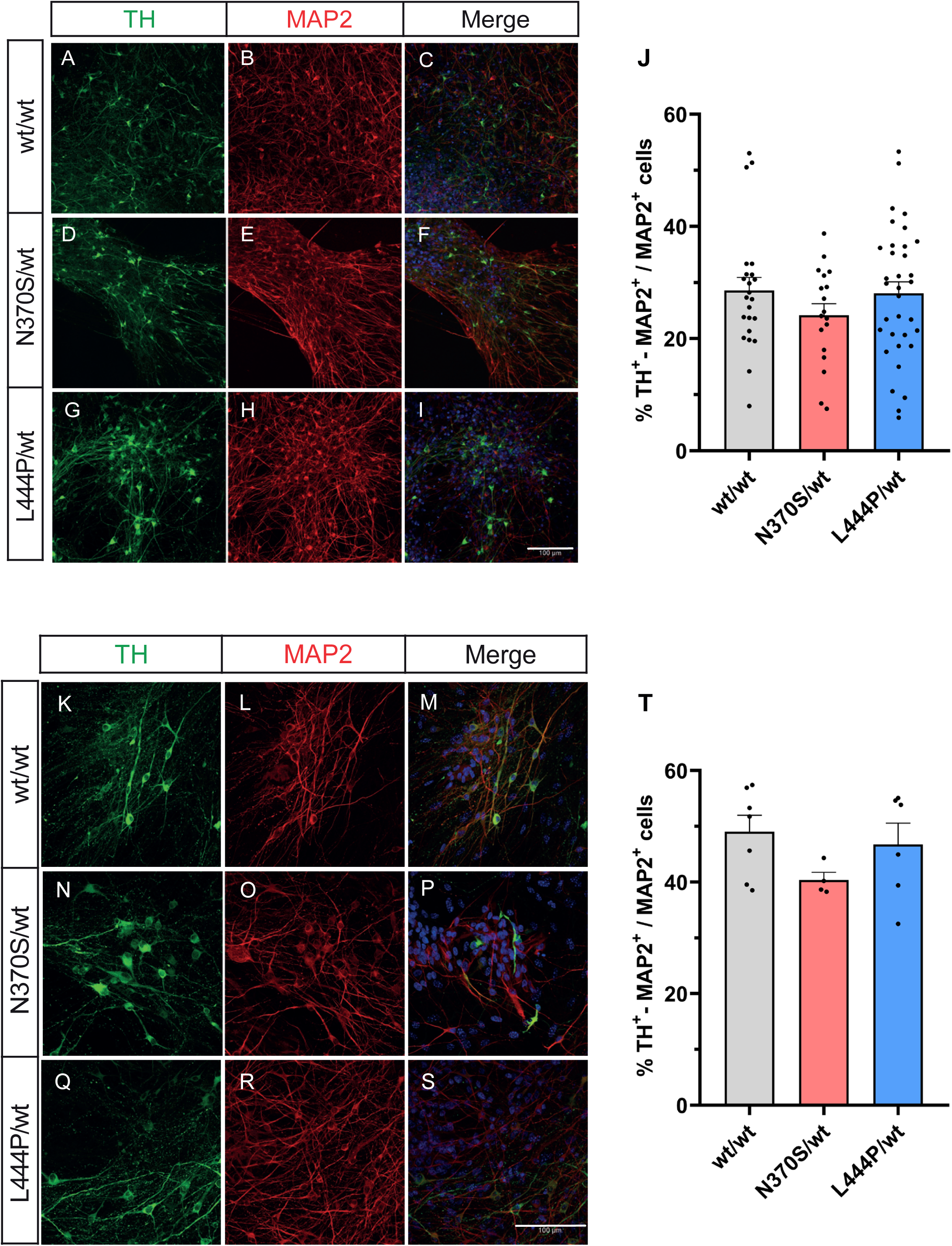
The N370S and L444P *GBA1* mutations do not significantly alter the proportion of iPSC-derived DA neurons. Cells differentiating from human iPSCs were fixed at 50 DIV (A-J) and 90 DIV (K-T), and then immunostained with specific antibodies against TH and MAP2. The percentage of TH^+^/MAP2^+^ neurons in the cultures was 24.0-28.2% at 50 DIV (J) and 40.4-49.0% at 90 DIV (T), and this parameter was not affected by either mutation. The results are the mean (±S.E.M.) of *n* = 18-33 cultures per genotype from 5-6 experiments of 50 DIV cultures, or of *n* = 4-7 cultures per genotype for 90 DIV cultures. Scale bars: 100 µm.

**Figure EV3.**
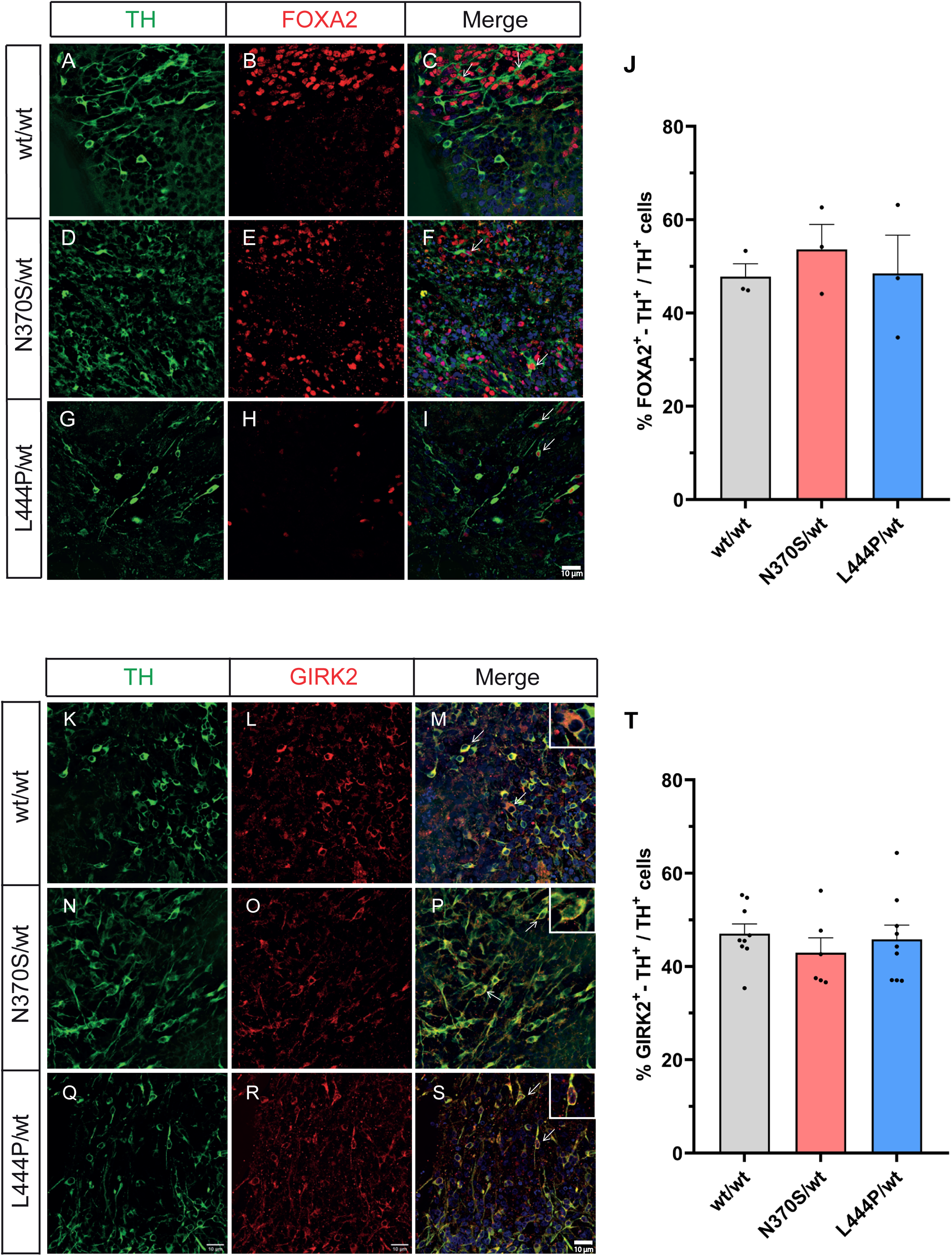
Expression of FOXA2 and GIRK2 in iPSC-derived DA neurons from *GBA1-*PD patients and healthy subjects. Neurons differentiating from human iPSCs were fixed at 21 DIV and immunostained with antibodies against TH and FOXA2 (A-J), or they were fixed at 54 DIV and immunostained with antibodies against TH and GIRK2 (K-T). The arrows point to double positive cells, some of which are presented at high magnification in the insets. Of the TH^+^ neurons, 47-53% also expressed FOXA2 (J) and 43.0-47.0% also expressed GIRK2 (T), with no effect of the mutations on these percentages. The results are the mean (±S.E.M.) of *n* = 3 cultures per genotype from 2-3 experiments stained with the TH and FOXA2 antibodies, or *n* = 6-9 cultures per genotype from 2-3 experiments stained with TH and GIRK2 antibodies. Scale bars: 10 µm.

**Figure EV4.**
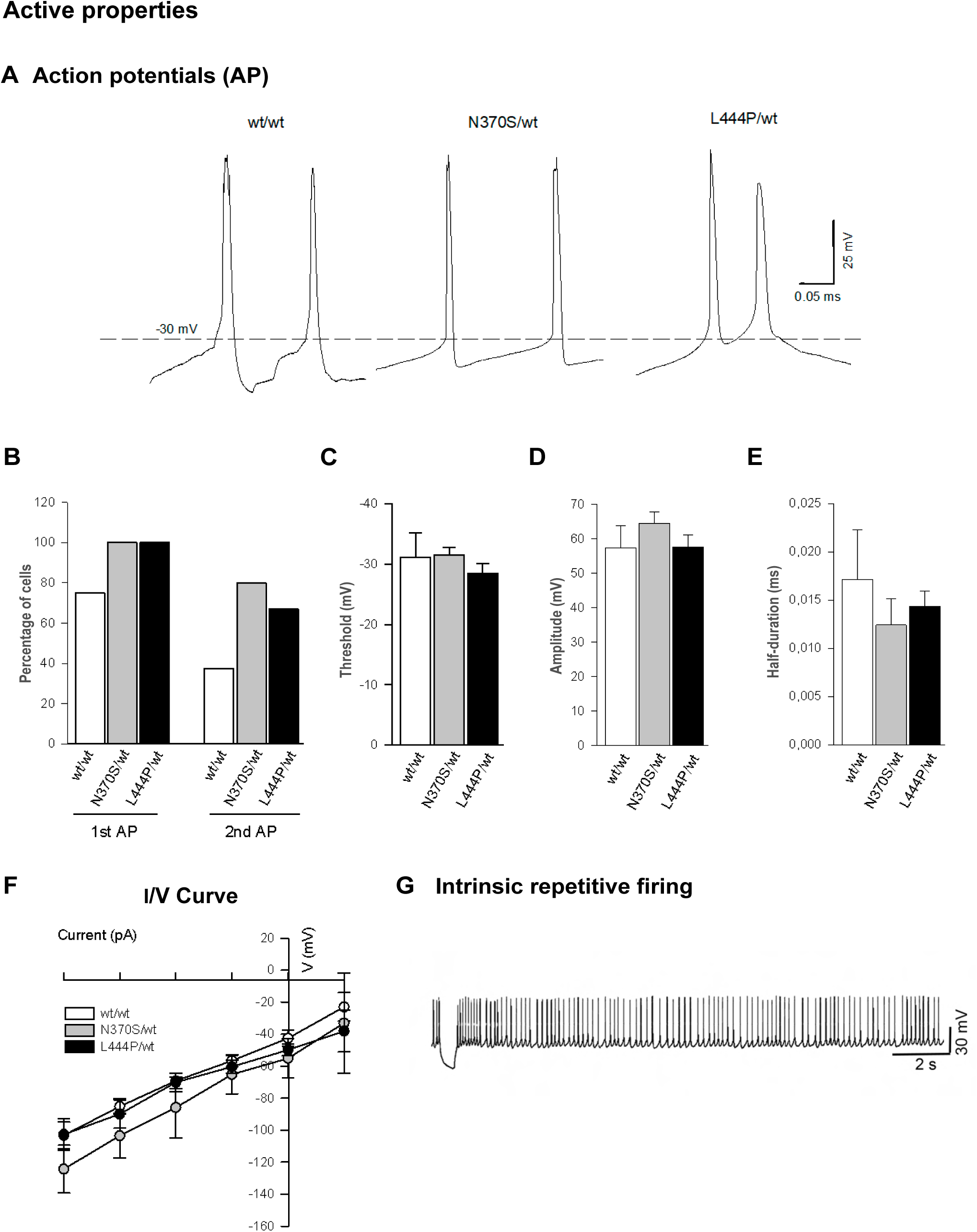
The active electrical properties of iPSC-derived DA neurons are little affected by the *GBA1* mutations. Neurons obtained from the human iPSCs of control subjects and PD patients carrying the N370S/wt or L444P/wt *GBA1* mutation were maintained for 90-93 days to perform electrophysiological recordings. The recorded neurons were injected with Alexa 488 and their dopaminergic phenotype was confirmed by immunostaining for TH (see supplementary Fig. S3A-F and S4A-L). The traces illustrating action potentials (APs) are shown in (A). The *GBA1* mutations increased, albeit not significantly, the proportion of neurons firing APs (B) and produced no changes in the AP characteristics (C-E) or in the I/V curve (F). G) An example of intrinsic repetitive firing of APs is shown. The results are the mean (±S.E.M.) of *n* = 6-12 neurons per genotype from 2 experiments.

## Reference

Alam P, Bousset L, Melki R, Otzen DE (2019) alpha-synuclein oligomers and fibrils: a spectrum of species, a spectrum of toxicities. J Neurochem 150: 522–534

Andersson E, Tryggvason U, Deng Q, Friling S, Alekseenko Z, Robert B, Perlmann T, Ericson J (2006) Identification of intrinsic determinants of midbrain dopamine neurons. Cell 124: 393–405

Baden P, Yu C, Deleidi M (2019) Insights into GBA Parkinson’s disease pathology and therapy with induced pluripotent stem cell model systems. Neurobiol Dis 127: 1–12

Beccano-Kelly DA, Cherubini M, Mousba Y, Cramb KML, Giussani S, Caiazza MC, Rai P, Vingill S, Bengoa-Vergniory N, Ng B et al (2023) Calcium dysregulation combined with mitochondrial failure and electrophysiological maturity converge in Parkinson’s iPSC-dopamine neurons. iScience 26: 107044

Blandini F, Cilia R, Cerri S, Pezzoli G, Schapira AHV, Mullin S, Lanciego JL (2019) Glucocerebrosidase mutations and synucleinopathies: Toward a model of precision medicine. Mov Disord 34: 9–21

Bloem BR, Okun MS, Klein C (2021) Parkinson’s disease. Lancet 397: 2284–2303

Brooker SM, Naylor GE, Krainc D (2024) Cell biology of Parkinson’s disease: Mechanisms of synaptic, lysosomal, and mitochondrial dysfunction. Curr Opin Neurobiol 85: 102841

Bruinsma IB, Bruggink KA, Kinast K, Versleijen AA, Segers-Nolten IM, Subramaniam V, Kuiperij HB, Boelens W, de Waal RM, Verbeek MM (2011) Inhibition of alpha-synuclein aggregation by small heat shock proteins. Proteins 79: 2956–2967

Buck SA, Erickson-Oberg MQ, Bhatte SH, McKellar CD, Ramanathan VP, Rubin SA, Freyberg Z (2022) Roles of VGLUT2 and Dopamine/Glutamate Co-Transmission in Selective Vulnerability to Dopamine Neurodegeneration. ACS Chem Neurosci 13: 187–193

Burbulla LF, Song P, Mazzulli JR, Zampese E, Wong YC, Jeon S, Santos DP, Blanz J, Obermaier CD, Strojny C et al (2017) Dopamine oxidation mediates mitochondrial and lysosomal dysfunction in Parkinson’s disease. Science 357: 1255–1261

Calabresi P, Di Lazzaro G, Marino G, Campanelli F, Ghiglieri V (2023) Advances in understanding the function of alpha-synuclein: implications for Parkinson’s disease. Brain 146: 3587–3597

Cavallieri F, Cury RG, Guimaraes T, Fioravanti V, Grisanti S, Rossi J, Monfrini E, Zedde M, Di Fonzo A, Valzania F et al (2023) Recent Advances in the Treatment of Genetic Forms of Parkinson’s Disease: Hype or Hope? Cells 12(5)

Chatterjee D, Krainc D (2023) Mechanisms of Glucocerebrosidase Dysfunction in Parkinson’s Disease. J Mol Biol 435: 168023

Chu Y, Kompoliti K, Cochran EJ, Mufson EJ, Kordower JH (2002) Age-related decreases in Nurr1 immunoreactivity in the human substantia nigra. J Comp Neurol 450: 203–214

Cilia R, Tunesi S, Marotta G, Cereda E, Siri C, Tesei S, Zecchinelli AL, Canesi M, Mariani CB, Meucci N et al (2016) Survival and dementia in GBA-associated Parkinson’s disease: The mutation matters. Ann Neurol 80: 662–673

Cox D, Carver JA, Ecroyd H (2014) Preventing alpha-synuclein aggregation: the role of the small heat-shock molecular chaperone proteins. Biochim Biophys Acta 1842: 1830–1843

Cramb KML, Beccano-Kelly D, Cragg SJ, Wade-Martins R (2023) Impaired dopamine release in Parkinson’s disease. Brain 146: 3117–3132

Doi D, Samata B, Katsukawa M, Kikuchi T, Morizane A, Ono Y, Sekiguchi K, Nakagawa M, Parmar M, Takahashi J (2014) Isolation of human induced pluripotent stem cell-derived dopaminergic progenitors by cell sorting for successful transplantation. Stem Cell Reports 2: 337–350

Dumas S, Wallen-Mackenzie A (2019) Developmental Co-expression of Vglut2 and Nurr1 in a Mes-Di-Encephalic Continuum Preceeds Dopamine and Glutamate Neuron Specification. Front Cell Dev Biol 7: 307

Eskenazi D, Malave L, Mingote S, Yetnikoff L, Ztaou S, Velicu V, Rayport S, Chuhma N (2021) Dopamine Neurons That Cotransmit Glutamate, From Synapses to Circuits to Behavior. Front Neural Circuits 15: 665386

Fares MB, Jagannath S, Lashuel HA (2021) Reverse engineering Lewy bodies: how far have we come and how far can we go? Nat Rev Neurosci 22: 111–131

Fernandes HJ, Hartfield EM, Christian HC, Emmanoulidou E, Zheng Y, Booth H, Bogetofte H, Lang C, Ryan BJ, Sardi SP et al (2016) ER Stress and Autophagic Perturbations Lead to Elevated Extracellular alpha-Synuclein in GBA-N370S Parkinson’s iPSC-Derived Dopamine Neurons. Stem Cell Reports 6: 342–356

Friling S, Andersson E, Thompson LH, Jonsson ME, Hebsgaard JB, Nanou E, Alekseenko Z, Marklund U, Kjellander S, Volakakis N et al (2009) Efficient production of mesencephalic dopamine neurons by Lmx1a expression in embryonic stem cells. Proc Natl Acad Sci U S A 106: 7613–7618

Garcia-Sanz P J MFGA, Moratalla R (2021) The Role of Cholesterol in alpha-Synuclein and Lewy Body Pathology in GBA1 Parkinson’s Disease. Mov Disord 36: 1070–1085

Garcia-Sanz P, Orgaz L, Bueno-Gil G, Espadas I, Rodriguez-Traver E, Kulisevsky J, Gutierrez A, Davila JC, Gonzalez-Polo RA, Fuentes JM et al (2017) N370S-GBA1 mutation causes lysosomal cholesterol accumulation in Parkinson’s disease. Mov Disord 32: 1409–1422

Garcia-Sanz P, Orgaz L, Fuentes JM, Vicario C, Moratalla R (2018) Cholesterol and multilamellar bodies: Lysosomal dysfunction in GBA-Parkinson disease. Autophagy 14: 717–718

Garritsen O, van Battum EY, Grossouw LM, Pasterkamp RJ (2023) Development, wiring and function of dopamine neuron subtypes. Nat Rev Neurosci 24: 134–152

Gegg ME, Menozzi E, Schapira AHV (2022) Glucocerebrosidase-associated Parkinson disease: Pathogenic mechanisms and potential drug treatments. Neurobiol Dis 166: 105663

Goker-Alpan O, Stubblefield BK, Giasson BI, Sidransky E (2010) Glucocerebrosidase is present in alpha-synuclein inclusions in Lewy body disorders. Acta Neuropathol 120: 641–649

Guzman JN, Sanchez-Padilla J, Chan CS, Surmeier DJ (2009) Robust pacemaking in substantia nigra dopaminergic neurons. J Neurosci 29: 11011–11019

Hayashi J, Carver JA (2020) The multifaceted nature of alphaB-crystallin. Cell Stress Chaperones 25: 639–654

Henderson MX, Sedor S, McGeary I, Cornblath EJ, Peng C, Riddle DM, Li HL, Zhang B, Brown HJ, Olufemi MF et al (2020) Glucocerebrosidase Activity Modulates Neuronal Susceptibility to Pathological alpha-Synuclein Insult. Neuron 105: 822–836 e827

Hnasko TS, Chuhma N, Zhang H, Goh GY, Sulzer D, Palmiter RD, Rayport S, Edwards RH (2010) Vesicular glutamate transport promotes dopamine storage and glutamate corelease in vivo. Neuron 65: 643–656

Horowitz M, Braunstein H, Zimran A, Revel-Vilk S, Goker-Alpan O (2022) Lysosomal functions and dysfunctions: Molecular and cellular mechanisms underlying Gaucher disease and its association with Parkinson disease. Adv Drug Deliv Rev 187: 114402

Jiang Z, Huang Y, Zhang P, Han C, Lu Y, Mo Z, Zhang Z, Li X, Zhao S, Cai F et al (2020) Characterization of a pathogenic variant in GBA for Parkinson’s disease with mild cognitive impairment patients. Mol Brain 13: 102

Kamath T, Abdulraouf A, Burris SJ, Langlieb J, Gazestani V, Nadaf NM, Balderrama K, Vanderburg C, Macosko EZ (2022) Single-cell genomic profiling of human dopamine neurons identifies a population that selectively degenerates in Parkinson’s disease. Nat Neurosci 25: 588–595

Kashani A, Betancur C, Giros B, Hirsch E, El Mestikawy S (2007) Altered expression of vesicular glutamate transporters VGLUT1 and VGLUT2 in Parkinson disease. Neurobiol Aging 28: 568–578

Kirkeby A, Grealish S, Wolf DA, Nelander J, Wood J, Lundblad M, Lindvall O, Parmar M (2012) Generation of regionally specified neural progenitors and functional neurons from human embryonic stem cells under defined conditions. Cell Rep 1: 703–714

Kittappa R, Chang WW, Awatramani RB, McKay RD (2007) The foxa2 gene controls the birth and spontaneous degeneration of dopamine neurons in old age. PLoS Biol 5: e325

Kouwenhoven WM, Fortin G, Penttinen AM, Florence C, Delignat-Lavaud B, Bourque MJ, Trimbuch T, Luppi MP, Salvail-Lacoste A, Legault P et al (2020) VGluT2 Expression in Dopamine Neurons Contributes to Postlesional Striatal Reinnervation. J Neurosci 40: 8262–8275

Kriks S, Shim JW, Piao J, Ganat YM, Wakeman DR, Xie Z, Carrillo-Reid L, Auyeung G, Antonacci C, Buch A et al (2011) Dopamine neurons derived from human ES cells efficiently engraft in animal models of Parkinson’s disease. Nature 480: 547–551

Kuo SH, Tasset I, Cheng MM, Diaz A, Pan MK, Lieberman OJ, Hutten SJ, Alcalay RN, Kim S, Ximenez-Embun P et al (2022) Mutant glucocerebrosidase impairs alpha-synuclein degradation by blockade of chaperone-mediated autophagy. Sci Adv 8: eabm6393

Kutzing MK, Langhammer CG, Luo V, Lakdawala H, Firestein BL (2010) Automated Sholl analysis of digitized neuronal morphology at multiple scales. J Vis Exp Nov 14: 45

Lang C, Campbell KR, Ryan BJ, Carling P, Attar M, Vowles J, Perestenko OV, Bowden R, Baig F, Kasten M et al (2019) Single-Cell Sequencing of iPSC-Dopamine Neurons Reconstructs Disease Progression and Identifies HDAC4 as a Regulator of Parkinson Cell Phenotypes. Cell Stem Cell 24: 93–106 e106

Lee HS, Bae EJ, Yi SH, Shim JW, Jo AY, Kang JS, Yoon EH, Rhee YH, Park CH, Koh HC et al (2010) Foxa2 and Nurr1 synergistically yield A9 nigral dopamine neurons exhibiting improved differentiation, function, and cell survival. Stem Cells 28: 501–512

Li H, Jiang H, Li H, Li L, Yan Z, Feng J (2022) Generation of human A9 dopaminergic pacemakers from induced pluripotent stem cells. Mol Psychiatry 27: 4407–4418

Lindersson E, Beedholm R, Hojrup P, Moos T, Gai W, Hendil KB, Jensen PH (2004) Proteasomal inhibition by alpha-synuclein filaments and oligomers. J Biol Chem 279: 12924–12934

Liu Y, Zhou Q, Tang M, Fu N, Shao W, Zhang S, Yin Y, Zeng R, Wang X, Hu G et al (2015) Upregulation of alphaB-crystallin expression in the substantia nigra of patients with Parkinson’s disease. Neurobiol Aging 36: 1686–1691

Masilamoni JG, Jesudason EP, Baben B, Jebaraj CE, Dhandayuthapani S, Jayakumar R (2006) Molecular chaperone alpha-crystallin prevents detrimental effects of neuroinflammation. Biochim Biophys Acta 1762: 284–293

Mata IF, Leverenz JB, Weintraub D, Trojanowski JQ, Chen-Plotkin A, Van Deerlin VM, Ritz B, Rausch R, Factor SA, Wood-Siverio C et al (2016) GBA Variants are associated with a distinct pattern of cognitive deficits in Parkinson’s disease. Mov Disord 31: 95–102

Mazzulli JR, Xu YH, Sun Y, Knight AL, McLean PJ, Caldwell GA, Sidransky E, Grabowski GA, Krainc D (2011) Gaucher disease glucocerebrosidase and alpha-synuclein form a bidirectional pathogenic loop in synucleinopathies. Cell 146: 37–52

Mazzulli JR, Zunke F, Isacson O, Studer L, Krainc D (2016) alpha-Synuclein-induced lysosomal dysfunction occurs through disruptions in protein trafficking in human midbrain synucleinopathy models. Proc Natl Acad Sci U S A 113: 1931–1936

Mendez I, Sanchez-Pernaute R, Cooper O, Vinuela A, Ferrari D, Bjorklund L, Dagher A, Isacson O (2005) Cell type analysis of functional fetal dopamine cell suspension transplants in the striatum and substantia nigra of patients with Parkinson’s disease. Brain 128: 1498–1510

Nieto-Estevez V, Pignatelli J, Arauzo-Bravo MJ, Hurtado-Chong A, Vicario-Abejon C (2013) A global transcriptome analysis reveals molecular hallmarks of neural stem cell death, survival, and differentiation in response to partial FGF-2 and EGF deprivation. PLoS One 8: e53594

Obeso JA, Monje MHG, Matarazzo M (2022) Major advances in Parkinson’s disease over the past two decades and future research directions. Lancet Neurol 21: 1076–1079

Onal G, Yalcin-Cakmakli G, Ozcelik CE, Boussaad I, Seker UOS, Fernandes HJR, Demir H, Kruger R, Elibol B, Dokmeci S et al (2024) Variant-specific effects of GBA1 mutations on dopaminergic neuron proteostasis. J Neurochem Apr 20

Patnaik S, Zheng W, Choi JH, Motabar O, Southall N, Westbroek W, Lea WA, Velayati A, Goldin E, Sidransky E et al (2012) Discovery, structure-activity relationship, and biological evaluation of noninhibitory small molecule chaperones of glucocerebrosidase. J Med Chem 55: 5734–5748

Pereira Luppi M, Azcorra M, Caronia-Brown G, Poulin JF, Gaertner Z, Gatica S, Moreno-Ramos OA, Nouri N, Dubois M, Ma YC et al (2021) Sox6 expression distinguishes dorsally and ventrally biased dopamine neurons in the substantia nigra with distinctive properties and embryonic origins. Cell Rep 37: 109975

Pfaffl MW (2001) A new mathematical model for relative quantification in real-time RT-PCR. Nucleic Acids Res 29: e45

Poewe W, Seppi K, Marini K, Mahlknecht P (2020) New hopes for disease modification in Parkinson’s Disease. Neuropharmacology 171: 108085

Pradas E, Martinez-Vicente M (2023) The Consequences of GBA Deficiency in the Autophagy-Lysosome System in Parkinson’s Disease Associated with GBA. Cells 12

Pu J, Lin L, Jiang H, Hu Z, Li H, Yan Z, Zhang B, Feng J (2023) Parkin Maintains Robust Pacemaking in Human Induced Pluripotent Stem Cell-Derived A9 Dopaminergic Neurons. Mov Disord 38: 1273–1281

Reyes S, Fu Y, Double K, Thompson L, Kirik D, Paxinos G, Halliday GM (2012) GIRK2 expression in dopamine neurons of the substantia nigra and ventral tegmental area. J Comp Neurol 520: 2591–2607

Rodriguez-Traver E, Diaz-Guerra E, Rodriguez C, Arenas F, Orera M, Kulisevsky J, Moratalla R, Vicario C (2020) A collection of three integration-free iPSCs derived from old male and female healthy subjects. Stem Cell Res 42: 101663

Rodriguez-Traver E, Diaz-Guerra E, Rodriguez C, Fernandez P, Arenas F, Arauzo-Bravo M, Orera M, Kulisevsky J, Moratalla R, Vicario C (2019a) A collection of integration-free iPSCs derived from Parkinson’s disease patients carrying mutations in the GBA1 gene. Stem Cell Res 38: 101482

Rodriguez-Traver E, Rodriguez C, Diaz-Guerra E, Arenas F, Arauzo-Bravo M, Orera M, Kulisevsky J, Moratalla R, Vicario C (2019b) Generation of an integration-free iPSC line, ICCSICi005-A, derived from a Parkinson’s disease patient carrying the L444P mutation in the GBA1 gene. Stem Cell Res 40: 101578

Rodriguez-Traver E, Solis O, Diaz-Guerra E, Ortiz O, Vergano-Vera E, Mendez-Gomez HR, Garcia-Sanz P, Moratalla R, Vicario-Abejon C (2016) Role of Nurr1 in the Generation and Differentiation of Dopaminergic Neurons from Stem Cells. Neurotox Res 30: 14–31

Rodriguez M, Gonzalez-Hernandez T (1999) Electrophysiological and morphological evidence for a GABAergic nigrostriatal pathway. J Neurosci 19: 4682–4694

Rosety I, Zagare A, Saraiva C, Nickels S, Antony P, Almeida C, Glaab E, Halder R, Velychko S, Rauen T et al (2023) Impaired neuron differentiation in GBA-associated Parkinson’s disease is linked to cell cycle defects in organoids. NPJ Parkinsons Dis 9: 166

Sanchez-Danes A, Richaud-Patin Y, Carballo-Carbajal I, Jimenez-Delgado S, Caig C, Mora S, Di Guglielmo C, Ezquerra M, Patel B, Giralt A et al (2012) Disease-specific phenotypes in dopamine neurons from human iPS-based models of genetic and sporadic Parkinson’s disease. EMBO Mol Med 4: 380–395

Schapira AHV, Chaudhuri KR, Jenner P (2017) Non-motor features of Parkinson disease. Nat Rev Neurosci 18: 509

Scheidt T, Carozza JA, Kolbe CC, Aprile FA, Tkachenko O, Bellaiche MMJ, Meisl G, Peter QAE, Herling TW, Ness S et al (2021) The binding of the small heat-shock protein alphaB-crystallin to fibrils of alpha-synuclein is driven by entropic forces. Proc Natl Acad Sci U S A 118

Schmittgen TD, Livak KJ (2008) Analyzing real-time PCR data by the comparative C(T) method. Nat Protoc 3: 1101–1108

Schondorf DC, Aureli M, McAllister FE, Hindley CJ, Mayer F, Schmid B, Sardi SP, Valsecchi M, Hoffmann S, Schwarz LK et al (2014) iPSC-derived neurons from GBA1-associated Parkinson’s disease patients show autophagic defects and impaired calcium homeostasis. Nat Commun 5: 4028

Schondorf DC, Ivanyuk D, Baden P, Sanchez-Martinez A, De Cicco S, Yu C, Giunta I, Schwarz LK, Di Napoli G, Panagiotakopoulou V et al (2018) The NAD+ Precursor Nicotinamide Riboside Rescues Mitochondrial Defects and Neuronal Loss in iPSC and Fly Models of Parkinson’s Disease. Cell Rep 23: 2976–2988

Seto-Salvia N, Pagonabarraga J, Houlden H, Pascual-Sedano B, Dols-Icardo O, Tucci A, Paisan-Ruiz C, Campolongo A, Anton-Aguirre S, Martin I et al (2012) Glucocerebrosidase mutations confer a greater risk of dementia during Parkinson’s disease course. Mov Disord 27: 393–399

Shahmoradian SH, Lewis AJ, Genoud C, Hench J, Moors TE, Navarro PP, Castano-Diez D, Schweighauser G, Graff-Meyer A, Goldie KN et al (2019) Lewy pathology in Parkinson’s disease consists of crowded organelles and lipid membranes. Nat Neurosci 22: 1099–1109

Shen H, Marino RAM, McDevitt RA, Bi GH, Chen K, Madeo G, Lee PT, Liang Y, De Biase LM, Su TP et al (2018) Genetic deletion of vesicular glutamate transporter in dopamine neurons increases vulnerability to MPTP-induced neurotoxicity in mice. Proc Natl Acad Sci U S A 115: E11532–E11541

Shults CW (2006) Lewy bodies. Proc Natl Acad Sci U S A 103: 1661–1668

Sidransky E, Lopez G (2012) The link between the GBA gene and parkinsonism. Lancet Neurol 11: 986–998

Sidransky E, Nalls MA, Aasly JO, Aharon-Peretz J, Annesi G, Barbosa ER, Bar-Shira A, Berg D, Bras J, Brice A et al (2009) Multicenter analysis of glucocerebrosidase mutations in Parkinson’s disease. N Engl J Med 361: 1651–1661

Song P, Peng W, Sauve V, Fakih R, Xie Z, Ysselstein D, Krainc T, Wong YC, Mencacci NE, Savas JN et al (2023) Parkinson’s disease-linked parkin mutation disrupts recycling of synaptic vesicles in human dopaminergic neurons. Neuron 111: 3775–3788 e3777

Steinkellner T, Conrad WS, Kovacs I, Rissman RA, Lee EB, Trojanowski JQ, Freyberg Z, Roy S, Luk KC, Lee VM et al (2021) Dopamine neurons exhibit emergent glutamatergic identity in Parkinson’s disease. Brain Nov 19

Steinkellner T, Zell V, Farino ZJ, Sonders MS, Villeneuve M, Freyberg RJ, Przedborski S, Lu W, Freyberg Z, Hnasko TS (2018) Role for VGLUT2 in selective vulnerability of midbrain dopamine neurons. J Clin Invest 128: 774–788

Stojkovska I, Wani WY, Zunke F, Belur NR, Pavlenko EA, Mwenda N, Sharma K, Francelle L, Mazzulli JR (2021) Rescue of alpha-synuclein aggregation in Parkinson’s patient neurons by synergistic enhancement of ER proteostasis and protein trafficking. Neuron Nov 9

Subramaniam M, Kern B, Vogel S, Klose V, Schneider G, Roeper J (2014) Selective increase of in vivo firing frequencies in DA SN neurons after proteasome inhibition in the ventral midbrain. Eur J Neurosci 40: 2898–2909

Surmeier DJ, Obeso JA, Halliday GM (2017) Selective neuronal vulnerability in Parkinson disease. Nat Rev Neurosci 18: 101–113

Tomagra G, Franchino C, Cesano F, Chiarion G, de Iure A, Carbone E, Calabresi P, Mesin L, Picconi B, Marcantoni A et al (2023) Alpha-synuclein oligomers alter the spontaneous firing discharge of cultured midbrain neurons. Front Cell Neurosci 17: 1078550

Tozzi A, Sciaccaluga M, Loffredo V, Megaro A, Ledonne A, Cardinale A, Federici M, Bellingacci L, Paciotti S, Ferrari E et al (2021) Dopamine-dependent early synaptic and motor dysfunctions induced by alpha-synuclein in the nigrostriatal circuit. Brain 144: 3477–3491

Tritsch NX, Ding JB, Sabatini BL (2012) Dopaminergic neurons inhibit striatal output through non-canonical release of GABA. Nature 490: 262–266

Trudeau LE, Hnasko TS, Wallen-Mackenzie A, Morales M, Rayport S, Sulzer D (2014) The multilingual nature of dopamine neurons. Prog Brain Res 211: 141–164

Woodard CM, Campos BA, Kuo SH, Nirenberg MJ, Nestor MW, Zimmer M, Mosharov EV, Sulzer D, Zhou H, Paull D et al (2014) iPSC-derived dopamine neurons reveal differences between monozygotic twins discordant for Parkinson’s disease. Cell Rep 9: 1173–1182

Wulansari N, Darsono WHW, Woo HJ, Chang MY, Kim J, Bae EJ, Sun W, Lee JH, Cho IJ, Shin H et al (2021) Neurodevelopmental defects and neurodegenerative phenotypes in human brain organoids carrying Parkinson’s disease-linked DNAJC6 mutations. Sci Adv 7

Yap TL, Gruschus JM, Velayati A, Westbroek W, Goldin E, Moaven N, Sidransky E, Lee JC (2011) Alpha-synuclein interacts with Glucocerebrosidase providing a molecular link between Parkinson and Gaucher diseases. J Biol Chem 286: 28080–28088

Zalon AJ, Quiriconi DJ, Pitcairn C, Mazzulli JR (2024) alpha-Synuclein: Multiple pathogenic roles in trafficking and proteostasis pathways in Parkinson’s disease. Neuroscientist: 10738584241232963

Zhang S, Wang R, Wang G (2019) Impact of Dopamine Oxidation on Dopaminergic Neurodegeneration. ACS Chem Neurosci 10: 945–953

Zunke F, Moise AC, Belur NR, Gelyana E, Stojkovska I, Dzaferbegovic H, Toker NJ, Jeon S, Fredriksen K, Mazzulli JR (2018) Reversible Conformational Conversion of alpha-Synuclein into Toxic Assemblies by Glucosylceramide. Neuron 97: 92–107 e110

