## Supplementary material for "*GBA1* mutations alter neuronal firing and structure, regulating VGLUT2 and CRYAB in dopamine neurons": Complete Methods and Supplemental figures

##### Differentiation of human iPSCs (hiPSCs) into DA neurons

The vector-free hiPSCs used in this study were generated and characterized previously in our laboratory (Rodriguez-Traver et al, 2020; Rodriguez-Traver et al, 2019a; Rodriguez-Traver et al, 2019b) and they have been deposited in the Spanish National Stem Cell Bank/Banco Nacional de Líneas Celulares (BNLC, Instituto de Salud Carlos III, Spain).

To differentiate the hiPSCs into DA neurons, we designed a protocol based on previous publications (Doi et al, 2014; Kirkeby et al, 2012; Kriks et al, 2011). First, the hiPSC colonies grown on mouse embryonic fibroblasts (MEFs) were dissociated into a single cell suspension by incubating them with accutase for 15-25 minutes at 37 ºC in an atmosphere of 5% $CO_2$. As this treatment also dissociates the MEFs, the cell suspension recovered was plated on 0.1% gelatin-coated culture dishes and left for 1.5 hours. After this period the MEFs had adhered to the gelatin, the supernatant containing the hiPSCs could be collected and the cells recovered by centrifugation for 5 minutes at 200g. In a pilot study, this supernatant was then seeded on coated Thermanox plastic coverslips and the cells were immunostained with antibodies against NANOG to ensure the purity of the hiPSC preparation. For differentiation, the pellet of hiPSC was resuspended in MEF-conditioned hiPSC medium supplemented with FGF-2 (6 ng/mL) and Y-27632 (10 µM). The cell suspensions were plated on vitronectin-coated culture dishes at a density of 50,000-80,000 cells/cm$^2$, and the medium was changed every other day using MEF-conditioned hiPSC medium supplemented with FGF-2.

Once the cultures reached a cell confluence of 90-100%, neural induction commenced by changing the medium to hiPSC medium supplemented with Noggin (250 ng/mL, day 0), containing A83-01 (5 μM) or SB431542 (10 μM) in fewer experiments. The medium was changed on day 1 of the culture to hiPSC medium supplemented with Noggin, A83, Sonic Hedgehog C25II (SHH, 100 ng/mL: R&D Systems, RYD-464-SH-025), Purmorphamine (2 μM: Miltenyi Biotec, 130-104-465) and FGF-8 (100 ng/mL: Peprotech,100-25B). On day 3, CHIR99021 (CHIR, 3 μM: Miltenyi Biotec, 130-103-926) was also added along with the factors added on day 1. From day 5 to day 9, the hiPSC medium was incrementally shifted to DMEMF12N2 medium (25%, 50%, 75%) every 2 days, and on day 7 and 9 the medium was only supplemented with Noggin and CHIR. On day 11, the medium was changed to Neurobasal/B27/GlutaMax supplemented with CHIR and with BDNF (20 ng/mL), ascorbic acid (0.2 mM), GDNF (20 ng/mL), dibutyril cAMP (0.5 mM) and TGFβ3 (1 ng/mL: BAGCT compounds). From day 13, CHIR was no longer added, BAGCT was included and the medium was changed every other day until the cells were dissociated with accutase and collected on days 17-20 (depending on the confluence of the cultures). The cells recovered were then plated on coverslips in the wells of 24 multi-well plates, or on 48, 12 or 6 multi-well plates without coverslips or on a layer of mouse astrocytes (see below). Before plating, both the coverslips and wells were coated with different combinations of molecules (including poly-ornithine, poly-D-lysine, fibronectin, laminin or matrigel) to ensure the best attachment and the differentiation-maturation of neurons. No clear differences in the number of neurons were observed between the different coatings and therefore, all the results obtained were combined.

For long-term maturation studies, cells were plated on a layer of mitotically inactivated primary astrocytes obtained from the cerebral cortex of postnatal day 0-2 (P0-P2) C57Bl/6 mice (kindly donated by Drs I. Torres-Alemán, L.M. García-Segura, and M.A. Arévalo, Cajal Institute, Madrid, Spain). The astrocytes were plated at 50,000-65,000 cells/cm$^2$ on poly-lysine-coated coverslips in DMEM containing 10% foetal bovine serum (FBS) and when confluent they were treated with cytosine-β-D-arabinofuranoside (AraC, 10 μM) for 2-3 days. The AraC was then removed and washed out with fresh medium.

The hiPSC-derived cells were collected, as mentioned above, resuspended in Neurobasal/B27/GlutaMax with BAGCT, and plated at 300,000-400,000 cells/cm$^2$ on coated coverslips and wells or at 150,000 cells/cm$^2$ on astrocytes. Depending on the studies carried out, the cells were maintained in culture from 18 to 104 days, changing the medium containing BAGCT every 2-3 days. Control, N370S-*GBA1* and L444P-*GBA*1 hiPSCs were always differentiated in parallel, and similarly, all the assays and analyses were performed in parallel.

The protocol using Noggin and A83-01 (5 μM) or Noggin and SB431542 (10 μM) favoured the production of tyrosine hydroxylase (TH)$^+$ neurons and the relatively stronger expression of mRNAs encoding dopaminergic transcripts (such as *TH*, *FOXA2*, *NURR1*, *LMX1A* and *LMX1B*) than other combination tested, including: LDN193189 (100 nM) and SB431542 (10 μM) or LDN193189 (100 nM) and A83-01 (5 μM). The proportion of neurons obtained after 30-33 DIV when LDN/SB, or LDN/A83 were used was very similar: 34.8% (±2.3%) of (β-III-Tubulin$^+$ or TUJ1$^+$ cells) cells were TH$^+$ (n=13), while 16.6% (±1.1%) of the total number of cells were TUJ1$^+$ and 6.1% (±0.1%) were TH$^+$. By contrast, adding Noggin/A83 or Noggin/SB, increased the proportion of TUJ1$^+$ and TH$^+$

neurons: 56.7% (±2.8%) of the TUJ1$^+$ cells were TH$^+$, and 33.5% (±3.0%) of the total cell population were TUJ1$^+$ and 18.3% (±1.4%) were TH$^+$. Noggin/A83 or Noggin/SB in fewer experiments, together with SHH, Pur, FGF8 and CHIR99021, were the molecules chosen to induce hiPSC differentiation into a midbrain dopaminergic cell fate.

**Real-Time Quantitative Reverse Transcription Polymerase Chain Reaction (RT-qPCR)**

After removing the medium, the cells were lysed in RA1/β-mercaptoethanol buffer and the extract was collected and immediately frozen in liquid nitrogen and stored at -80 ºC. The NucleoSpin RNA extraction kit (Macherey-Nagel, Cat. No. 740955.50) was used to purify the RNA from the cells, according to the manufacturer's instructions. The concentration and quality of the RNA was determined using NanoDrop One (Thermo Scientific, ND-ONE-W): absorbance ratios A260/280 >1.8 and A260/230 >1.5.

Complementary DNA (cDNA) was obtained by reverse transcription of 250-500 ng of the initial RNA using the SuperScript III enzyme (Invitrogen), according to the manufacturer's instructions and our protocols (Nieto-Estevez et al, 2013). To determine the relative expression of different genes the RT-qPCR analysis was performed in triplicate using Power SYBR Green or Power SYBR Green Fast with the appropriate negative controls (one negative control containing water instead of cDNA and a second, in which the negative control for the reverse transcriptase reaction was added instead of cDNA). The primer pairs, designed with either Lasergene software or purchased directly (Supplementary Table S1), were added to the SYBR Green mix, which was then added to each well of a 96 well-plate for RT-qPCR. Subsequently, the cDNA was added to each well and the plate was inserted into one of two RT-qPCR thermocyclers: 7500 Fast Real-Time PCR and QuantStudio 3 (Applied Biosystems), which use 7500 Software v2.3 and software QuantStudioTM Design & Analysis, respectively. All the RT-qPCR cDNA products obtained were sequenced to verify that they corresponded to the expected fragments.

For quantification, the Ct value for each target gene was normalized to the Ct value for *GAPDH* using the comparative $C_T$ method with the equation $2^{-\Delta C_T}$ (Pfaffl, 2001; Schmittgen & Livak, 2008) This method was used for all the genes as the slope obtained after representing the $C_T$ of each gene relative to that for *GAPDH* was between 0.1 and -0.1, indicating the efficiency of the qPCR reaction was the same for the gene studied and for *GAPDH*. Changes in gene expression were compared between the N370S/wt and L444P/wt cultures, and also with the expression in the control (wt/wt) cultures, using the comparative $C_T$ method with the equation $2^{-\Delta\Delta C_T}$. Values were obtained as the fold change on a log2 scale, and then represented using violin plots after conversion to a linear scale. Violin plots represent the density, distribution and the median of the data obtained from three differentiation experiments performed in triplicate. Statistical analysis was performed using an unpaired Student's t test, applying Welch's correction when the F-test indicated that the variances of both groups differed significantly.

**Immunocytochemistry (ICC)**

Cells were fixed for 25 minutes with 4% paraformaldehyde (PFA) and preserved at 4 ºC in sodium azide (0.02% in phosphate-buffered saline - PBS). For immunocytochemistry (ICC), cells were first washed three times with PBS and then incubated for 1 hour at room temperature (RmT) in permeabilization/blocking buffer (0.1-0.4% Triton X-100/Normal goat or donkey serum/PBS), prior to probing them 20-22 hours with primary antibodies against: CRYAB (1:250: Abcam Cat# ab76467, RRID:AB_1523120); FOXA2 (1:100: Abcam Cat# ab108422, RRID:AB_11157157); GABA (1:2000: Sigma Cat# A2051, RRID:AB_2314459); GIRK2 (1:200: Abcam Cat# ab65096, RRID:AB_1139732); MAP2 (1:250: Sigma-Aldrich Cat# M1406, RRID:AB_477171 and 1:1000: Synaptic Systems Cat# 188004, RRID:AB_2138181); α-synuclein LB509 (1:800: Abcam Cat# ab27766, RRID:AB_727020); TH (1:200: Millipore Cat# MAB318, RRID:AB_2201528 and 1:200: Millipore Cat# AB152, RRID:AB_390204); β-III-tubulin (TUJ1, 1:1000: Covance Cat# MMS-435P, RRID:AB_2313773 and TUJ1, 1:300: Abcam Cat# ab18207, RRID:AB_444319); VGLUT2 (1:500: Alomone Cat# AGC-036, RRID:AB_2340950). Antibody binding was then detected with the appropriate secondary antibodies for 1 h at RmT, while the nuclei were stained with Hoechst. No specific signal was detected in the absence of the primary antibodies but in the presence of the secondary antibodies.

For quantification, confocal images from 5 or 10 random microscope fields were used per culture using a 63x objective or a 40x objective with a 1.5 zoom. Cells in the entire Z-stack were counted and the co-localization of specific markers was analysed in each individual Z-plane using ImageJ software (NIH, Bethesda, MD). The proportion of labelled cells was calculated relative to the total number of $MAP2^+$, $TUJ1^+$ or $TH^+$ neurons, or of the total Hoechst stained nuclei. The results were expressed as the mean ± S.E.M. of 3-33 cultures per genotype from 3-7 differentiation experiments. The specific *n* values for each ICC assay are given in the figure legends and the statistical analysis was carried out using one-way ANOVA followed by a Bonferrani or Tukey post-hoc test, or Kruskal-Wallis test followed by a Dunn post-hoc test.

**Analysis of α-synuclein aggregates in DA neurons**

The hiPSC-derived cells were fixed on DIV 95 or 103, treated for 1h with 0.1-0.2% Triton X-100/Normal goat serum(NGS)/PBS, and probed for 20-22 h at 4 ºC with antibodies against TH, MAP2 and α-syn. Cells were then washed with PBS and incubated with the appropriate secondary antibodies for 1h at RmT, washed and mounted with Mowiol after staining the nuclei with Hoechst. Confocal images (each containing a clearly defined $TH^+$ neuron) were obtained with a 63x objective using a 3.5 zoom. The neurons from wt/wt, N370S/wt and L444P/wt genotypes were analysed, and three images were taken from each neuron: one from the soma and two from two dendrites. The number of α-syn aggregates was semi-automatically counted using ImageJ. First, a threshold for detecting $TH^+$ staining was set to detect the neuron of interest which was then manually traced and its somatic and dendritic area measured. Next, a threshold for α-$syn^+$ staining was applied to quantify the number of aggregates. This allowed for an automatic detection of aggregates above a specific threshold. The aggregates and areas were determined blind to the genotype, and expressed as the number of aggregates per somatic area or per dendritic area (in $\mu m^2$). Statistical analysis was performed using the non-parametric Kruskal-Wallis test followed by Dunn´s multiple comparisons test.

**Dopamine release**

Dopamine in the culture medium collected at 47, 51 and 59 DIV was assessed, and the results were combined in the analysis. To measure the basal dopamine, the culture medium was replaced with fresh medium for 24 hours before it was collected. Subsequently, fresh medium with KCl (56 mM) was added to the cells and after 20 min it was collected to measure the dopamine release evoked, while the cells were fixed with 4% PFA for TH ICC. All the supernatants were collected in the presence of EDTA (1 mM: Sigma-Aldrich) and sodium metabisulfite (4 mM: Sigma-Aldrich) to prevent dopamine degradation, and they were kept at -80 ºC. Dopamine was extracted and measured using the Dopamine Research Elisa™ assay kit, according to the manufacturer's instructions (Labor Diagnostika Nord GmbH & Co. KG). The final colorimetric reaction was monitored at 450 nm and the samples were quantified by comparing their absorbance to a reference curve prepared from known dopamine concentrations (after subtracting the absorbance of the blank wells). The cross reactivity with other amines and with non-amine molecules in the assay was <0.007%-0.55% (according to the manufacturer´s information). Calculations were made following a non-linear regression for curve fitting using GraphPad Prism 5.0. Fixed cells were immunostained with an antibody against TH to determine the total number of $TH^+$ cells per well. For this, 10 fields per well (20x, 2-fold zoom or 10x, 4-fold zoom), were taken randomly following an established pattern. The total number of $TH^+$ cells per well was estimated as the number of $TH^+$ cells counted divided by 10 fields and then by the area of a field (to give the number of $TH^+$ neurons per $\mu m^2$), multiplied by the area of the well. The dopamine data was expressed as picograms (pg) of dopamine per $TH^+$

neuron, or as pg of dopamine per $TH^+$ neuron and per minute, as the mean (±S.E.M.) of 13-24 cultures per genotype and from 5 experiments. Statistical analysis was performed using one-way ANOVA with a Tukey post-hoc test.

**Quantification of glutamate in the culture medium**

Basal glutamate levels in the culture medium of iPSC-differentiating neurons (47 DIV) were determined using an enzymatic and colorimetric assay (Glutamate Assay Kit, Sigma Aldrich, MAK4004) after protein precipitation with 1M perchloric acid followed by neutralization with 2M KOH. After centrifugation, the supernatant was collected and glutamate was determined by measuring the absorbance at 450 nm in a spectrophotometer (FLUOstar OPTIMA; BMG, Labtech). The glutamate data was expressed as glutamate (nM)/min or as glutamate (pM/min)/$VGLUT2^+$ -$TH^+$ neurons, as the mean (±S.E.M.) of 5-8 cultures per genotype and from 2 experiments. Statistical analysis was performed using one-way ANOVA with a Tukey post-hoc test.

**Quantification of α-synuclein in the culture medium**

The hiPSC derived neurons were maintained in culture for 47 or 90 DIV, when the medium was collected to determine the α-syn levels using the SimpleStep ELISA kits (Abcam ab210973 and ab 260052) according to the manufacturer´s instructions. No α-syn was detected in the α-syn knockout cells (according to the manufacturer), reflecting the specificity of this assay. The final colorimetric reaction was monitored at 450 nm and the samples were quantified by comparing their optical density (OD, after subtracting the OD of the blank wells) to a reference curve prepared from known concentrations of α-syn. Calculations followed a near-linear regression for fitting using GraphPad Prism 5.0 software and fixed cells were immunostained with an antibody against TH to determine the total number of $TH^+$ cells per well, as described above. The data were expressed as picograms (pg/ml) of α-syn per $TH^+$ neuron and are the mean (±S.E.M.) of 16-19 cultures per genotype from 2 experiments. Statistical analysis was performed using one-way ANOVA with a Tukey post-hoc test.

**Electrophysiology**

Electrophysiological experiments were performed on hiPSC-differentiated neurons (90-93 DIV) cultured on a monolayer of mouse astrocytes growing on coverslips, which were transferred to a submersion-type recording chamber mounted on a DM6000FS Leica microscope. The chamber was continuously perfused (0.5 ml/min) with carbogen (95% $O_2$, 5% $CO_2$)-bubbled artificial cerebrospinal fluid (aCSF) at 36.5 ºC. The aCSF medium was prepared daily, with an osmolarity adjusted to 310 mOsm with sucrose and containing (in mM): 130 NaCl, 4 KCl, 2 $CaCl_2$, 1 $MgCl_2$, 10 HEPES, and 10 glucose [pH 7.4].

Whole-cell patch-clamp pipettes (borosilicate with filament, 1.5 mm outer diameter thick glass) were pulled on a Flamming/Brown micropipette puller P-97 (Sutter Instruments Co, Novato, CA) and had an initial resistance of 5–10 MΩ. The internal recording solution was adjusted to an osmolarity of 290 mOsm with sucrose and it contained (in mM): 130 Cs-gluconate, 10 NaCl, 2 $MgCl_2$, 0.2 EGTA, 1 NaATP, and 10 HEPES adjusted to pH 7.2. To visualize the cells, Alexa Fluor-488 hydrazide sodium salt (125 μM) was added to the internal solution. Patch-clamp recordings were obtained in the whole-cell configuration using Axoclamp 2-B (Molecular Devices, Sunnyvale, CA, USA) and digitized at

3 KHz using 1440A Digidata (Molecular Devices). The amplifier bridge circuit was adjusted to compensate for serial resistance and monitored throughout the experiment. Passive properties (resting membrane potential -RMP, membrane capacitance -Cm, and membrane resistance -Rm) were analysed using the membrane test function integrated in the pClamp10 software. Neuronal excitability was assessed in current clamp mode with a 500 ms somatic current injection and spontaneous synaptic activity was recorded holding the cells at -70 mV (voltage clamp). The data were stored and analysed on a digital system (pClamp v10.0, Molecular Devices). Results are presented as the mean (±S.E.M.) of 6-12 DA neurons (90-93 DIV) from two experiments. The data was analysed using one- or two-way ANOVA with a Dunn or Bonferroni post-test, respectively, or using the Chi-squared test, with $P<0.05$ considered significant.

**Morphological analysis of the cells recorded**

Neurons were injected with Alexa Fluor-488 while electrophysiological recordings were carried out (see above) and subsequently, they were fixed with 4% PFA and immunostained with antibodies against TH and MAP2. The samples were scanned using an AF 6000LX Leica microscope with a 20x objective to identify the Alexa Fluor 488 labelled cells. Once the recorded neurons had been recognized, they were imaged using a confocal microscope with a 40x objective and analysed using Fiji ImageJ software. The cells were first classified as $TH^+$ or $TH^-$ and subsequently, $TH^+$ neurons were traced using ImageJ Sholl analysis software using the Bonfire program in MATLAB (Kutzing et al, 2010). The average length of the processes, their total length and the number of processes was also quantified. However, not all neurons recorded could be found on the coverslips after the electrophysiology experiments, probably because they died or because dye filling failed. Moreover, some $TH^+$ neurons were not completely filled with Alexa-488 and therefore, they were removed from the analysis.

In the Sholl analysis, the number of intersections depends on two factors, the genotype and the distance from the soma, such that a two-way ANOVA analysis with a Bonferroni´s test was performed. A one-way ANOVA with a Tukey´s post-test was used to analyse the length and number of processes per neuron. The results are expressed as the mean (±S.E.M.) of 8-9 neurons per genotype from 3 experiments.

**Electron microscopy**

The hiPSC-differentiated neurons (90-104 DIV) cultured on a mouse astrocyte monolayer on coated thermanox plastic coverslips were examined by transmission electron microscopy (TEM). The cells were fixed in 4% PFA and 2% glutaraldehyde in 0.12M Phosphate Buffer (PB, pH 7.4), and then treated for 45 minutes at RT with 1% $OsO_4$ (Electron Microscopy Sciences, Hatfield, PA, USA) and 7% glucose in PB, after which they were stained with 1% uranyl acetate (Electron Microscopy Sciences) in maleate buffer (pH 4.5). The cells were then dehydrated in increasing concentrations of ethanol (50°, 70°, 96° and 100°), and washed twice for 10 minutes in 100° ethanol and $CuSO_4$ to complete dehydration. After clearing in propylene oxide (Fluka AG, Buch, Switzerland) and embedding in Durcupan, the Durcupan was polymerized for 24 h at 60 °C. The next day, the coverslips were placed on top of pre-made resin columns and the resin was polymerized at at 60 ºC for 24 h. Ultrathin sections (75–80 nm) were obtained and then studied under transmissionTEM.

Schmittgen TD, Livak KJ (2008) Analyzing real-time PCR data by the comparative C(T) method. *Nat Protoc* 3: 1101-1108

**Supplementary Figure S1. DA neurons are generated in similar percentages from the iPSCs of *GBA1*-PD patients and healthy subjects.**

(A-I) Cells differentiating from human iPSCs were fixed at 30 DIV and then immunostained with specific antibodies against TH and β-III-tubulin (TUJ1). The nuclei were stained with Hoechst and the arrows indicate double labelled neurons.

(J) The percentage of $TH^+/TUJ1^+$ neurons in the cultures was 47.1-56.0%, without no significant effect of the mutations in this assay. The results are the mean (±S.E.M.) of *n* = 5-6 cultures per genotype from 3-4 experiments.

Scale bar: 100 μm.

**Supplementary Figure S2. A small subpopulation of iPSC-derived DA neurons express GABA.**

(A-I) Neurons differentiating from human iPSCs were fixed at 54 DIV and immunostained with antibodies against TH and GABA. The insets show examples of $GABA^+/TH^+$ neurons (arrows) at a higher magnification and the nuclei were stained with Hoeschst.

(J) Of the $TH^+$ neurons, 5.7%-7.3% expressed GABA, and the N370S/wt or L444P/wt *GBA1* mutation did not affect these percentages. The results are the mean (±S.E.M.) of *n* = 6-9 cultures per genotype from 2-3 experiments.

Scale bar: 10 μm.

**Supplementary Figure S3. The passive electrical properties of iPSC-derived DA neurons are not affected by the *GBA1* mutations.**

Neurons obtained from the human iPSCs of control subjects and PD patients carrying the N370S/wt or L444P/wt *GBA1* mutations were maintained for 90-93 days to perform electrophysiological recordings.

(A-F) Recorded neurons were injected with Alexa 488 and their dopaminergic phenotype was confirmed by immunostaining for TH (see supplementary S4A-L). The traces illustrating the passive properties are shown in (G). The *GBA1* mutations produced no significant changes in the RMP (H), Cm (I) and Rm (J). The results are the mean (±S.E.M.) of *n* = 6-12 neurons per genotype from 2 experiments. RMP, resting membrane potential; Cm, capacitance; Rm, resistance. Scale bar: 43.7 μm.

**Supplementary Figure S4. The effect of *GBA1* mutations on DA neuron morphology (I).**

Human iPSC-derived DA neurons from control subjects and PD patients carrying a N370S/wt or L444P/wt mutation in *GBA1* were maintained in culture for 90-93 days (A-L) The recorded neurons were injected with Alexa 488 and their dopaminergic neuronal phenotype was confirmed by immunostaining for TH and MAP2 (arrows indicate triple labelled neurons).

(M-P) A Sholl analysis was performed and the graphs show the influence of the *GBA1* genotype on the number of intersections relative to the distance from the center of the soma. Significant increases in the number (No.) of intersections in the first 20-65 μm were observed in L444P/wt DA neurons. The results are the mean (±S.E.M.) of *n* = 8-9 neurons per genotype from 3 experiments: **P*<0.05; **P*<0.01; ****P*<0.001 (two-way ANOVA followed by a Bonferroni´s test).

Scale bar: 100 μm.

**Supplementary Figure S5. The effect of *GBA1* mutations on DA neuron morphology (II).**

Human iPSC-derived neurons from control subjects and PD patients carrying the N370S/wt or L444P/wt mutation in *GBA1* were maintained in culture for 90-93 days. The recorded neurons were injected with Alexa 488 and their dopaminergic phenotype was confirmed by immunostaining for TH and MAP2.

(A-C) The mutations did not alter the length or the number of processes ($MAP2^{+}$ dendrites). The results are the mean (±S.E.M.) of $n$ = 8-9 neurons per genotype from 3 experiments.

**Supplementary Figure S6. N370S and L444P *GBA1* mutations promote autophagosome formation in iPSC-derived neurons.**

Low-magnification micrographs of 103 DIV cultures containing neurons derived from control healthy subjects (A) and from patients carrying the N370S/wt *GBA1* mutation (B). Note that mutant cells accumulate autophagosome-like structures in their processes (B, arrows). Autophagosome-like vacuoles are better visualized in the high magnification images of cells carrying the N370S/wt (C) or L444P/wt *GBA1* mutation (D). Scale bars: A-B, 2 µm; C-D, 500 nm.

**Supplementary Figure S7. Gene expression in post-mortem tissue from PD patients and healthy subjects.**

Gene expression analysis by RT-qPCR in samples from the substantia nigra (A) and hippocampal cortex (B) reveals no significant changes in selected transcripts between PD patients and controls. The violin plots represent the density, distribution and median of the data obtained by studying samples from 4 controls and 4 PD patients carrying no *GBA1* mutations, performed in triplicate.

### Figure S1

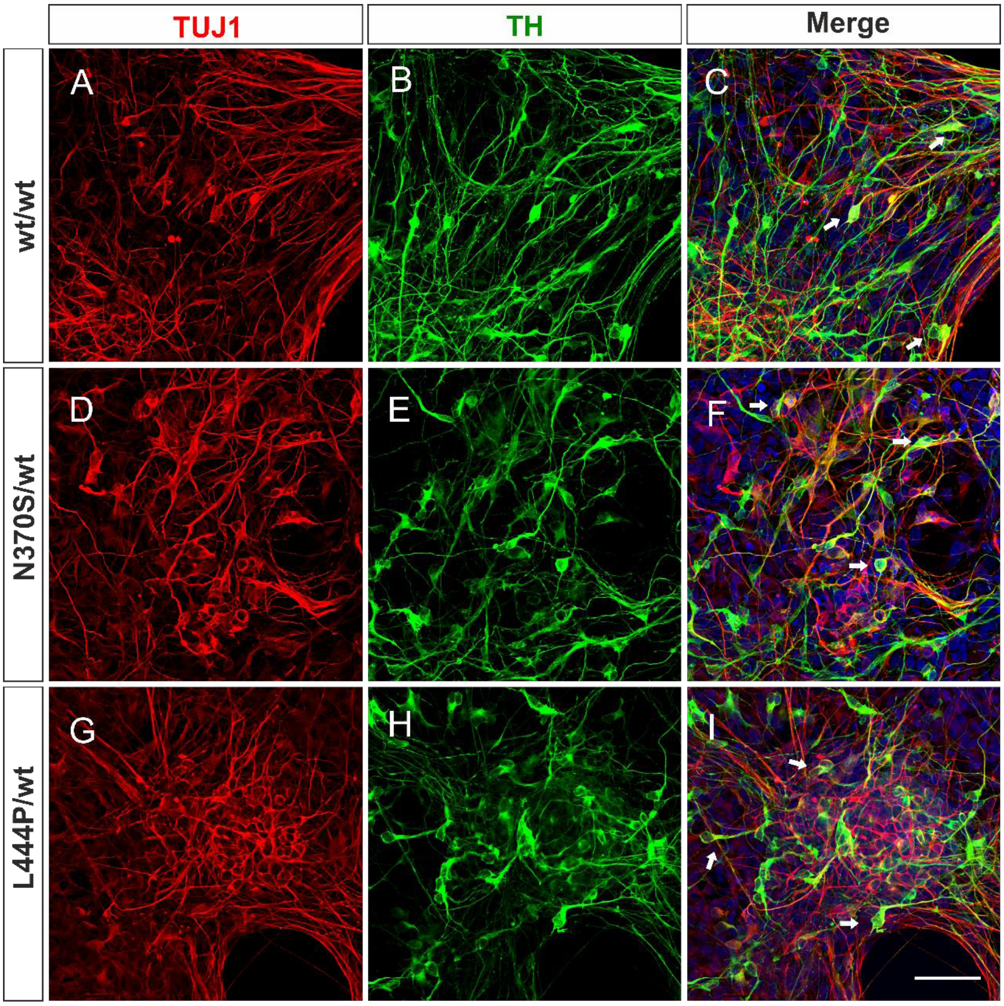

TUJ1
TH
Merge
wt/wt
N370S/wt
L444P/wt
A
B
C
D
E
F
G
H
I


**J**

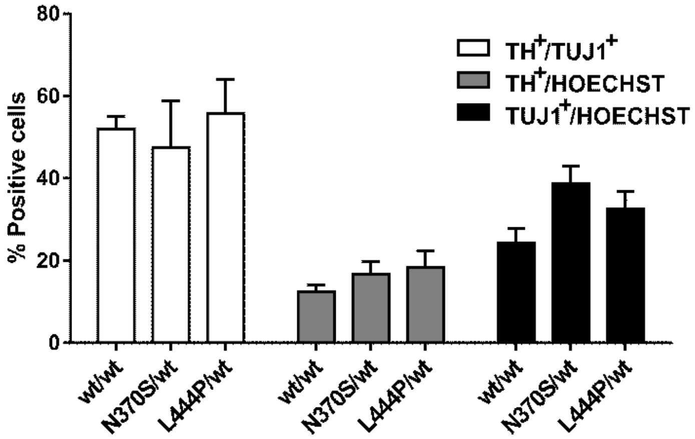

% Positive cells
80
60
40
20
0
TH+/TUJ1+
TH+/HOECHST
TUJ1+/HOECHST
wt/wt
N370S/wt
L444P/wt
wt/wt
N370S/wt
L444P/wt
wt/wt
N370S/wt
L444P/wt

**Figure S2**

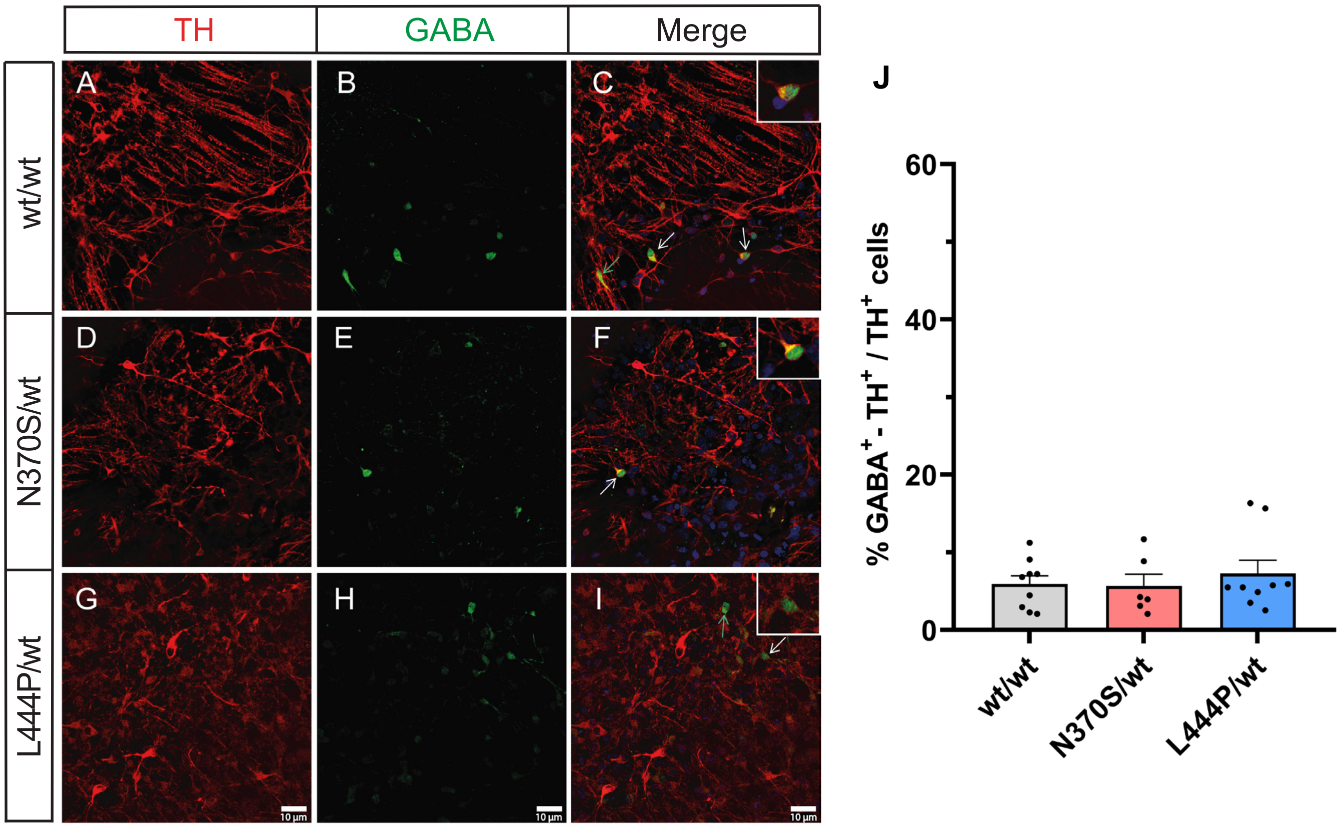
TH
GABA
Merge
wt/wt
N370S/wt
L444P/wt
A
B
C
D
E
F
G
H
I
10 µm
J
% GABA+ - TH+ / TH+ cells
60
40
20
0
wt/wt
N370S/wt
L444P/wt

**Figure S3**

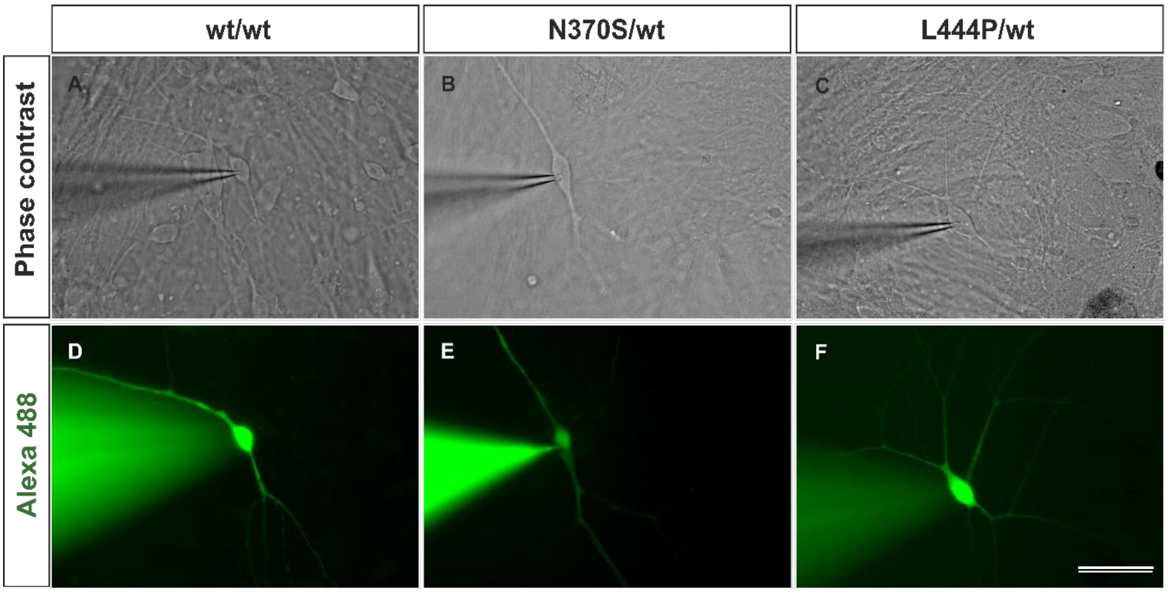


**Passive properties**

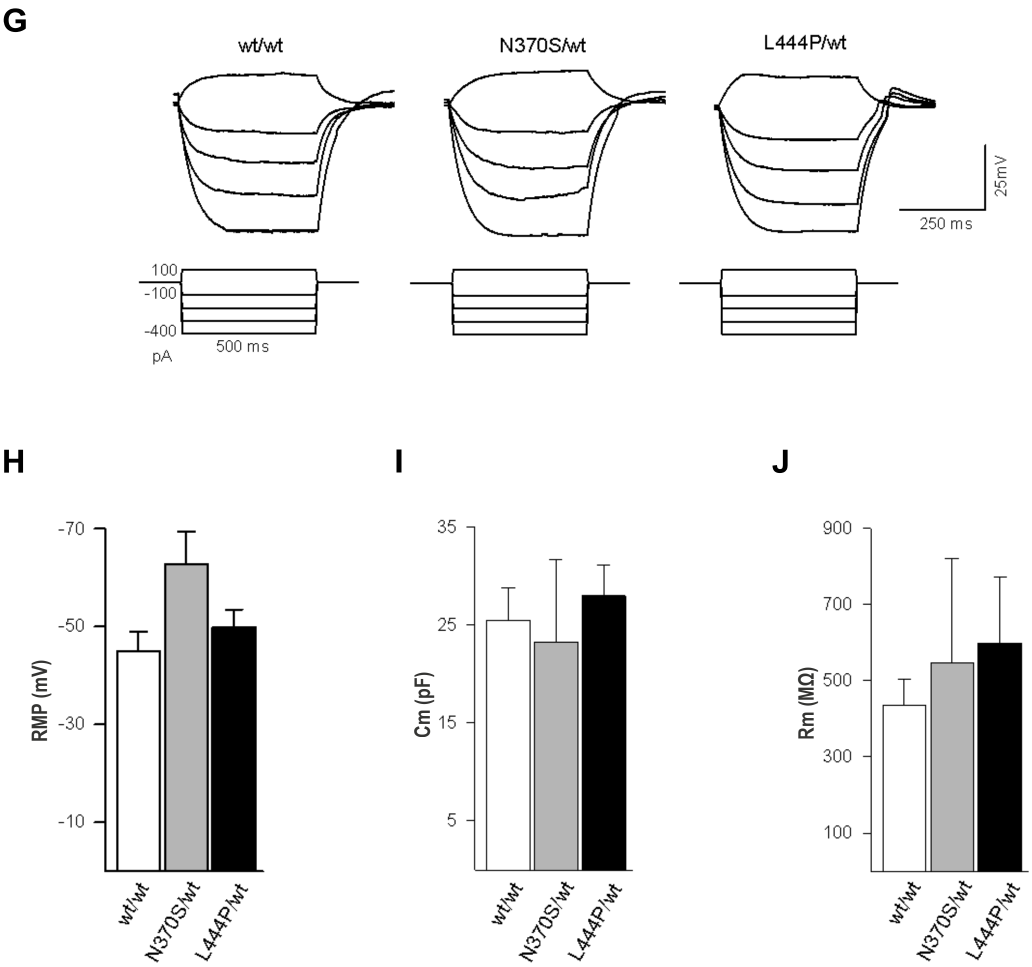

**Figure S4**

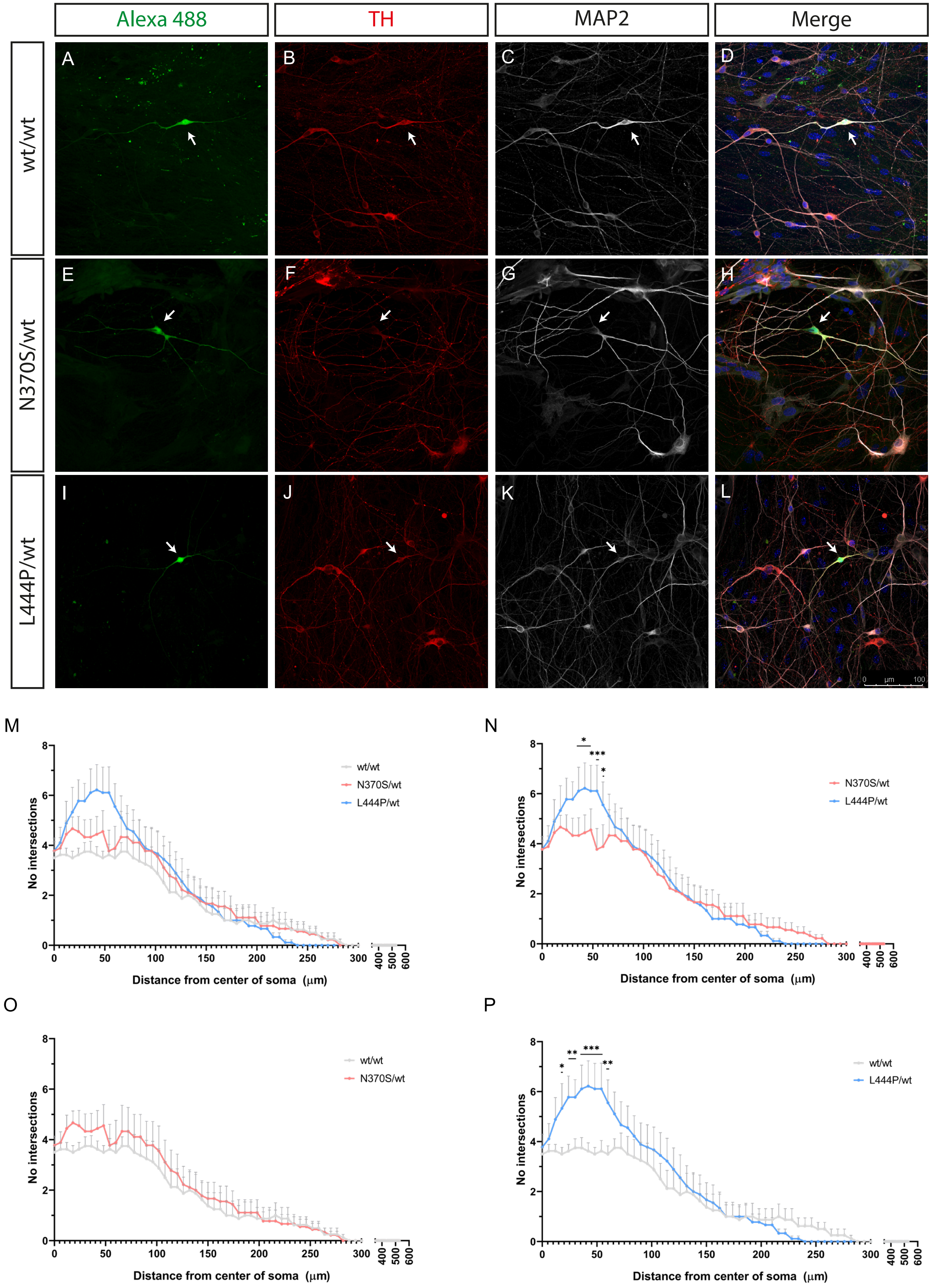

Alexa 488
TH
MAP2
Merge
wt/wt
N370S/wt
L444P/wt
A
B
C
D
E
F
G
H
I
J
K
L
0 μm 100
M
N
O
P
No intersections
Distance from center of soma (μm)
wt/wt
N370S/wt
L444P/wt
*
***
**

**Figure S5**

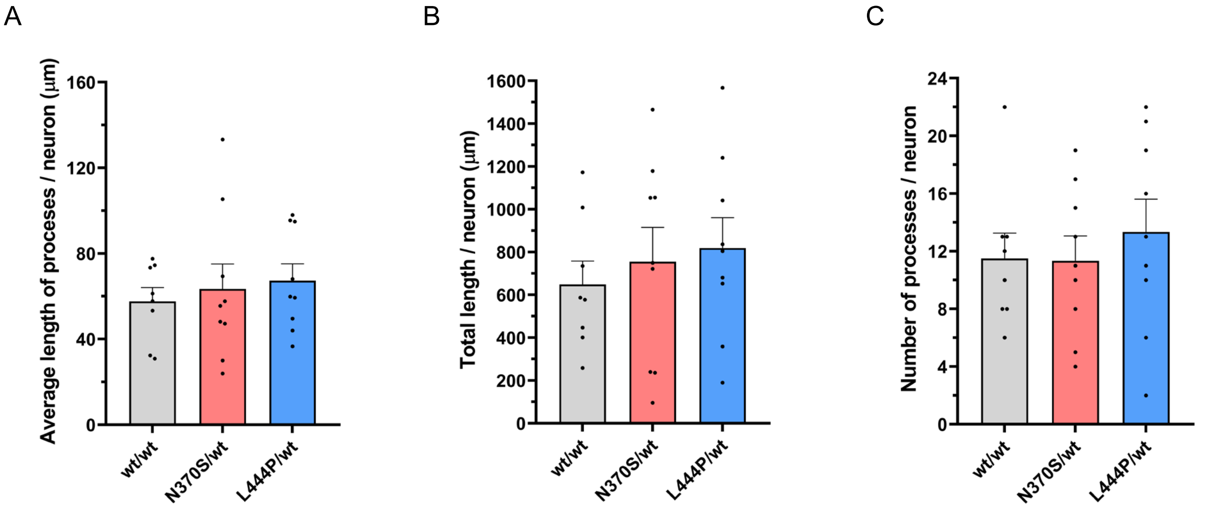
A
B
C
Average length of processes / neuron (μm)
Total length / neuron (μm)
Number of processes / neuron
160
120
80
40
0
1600
1400
1200
1000
800
600
400
200
24
20
16
12
8
4
wt/wt
N370S/wt
L444P/wt

Figure S6

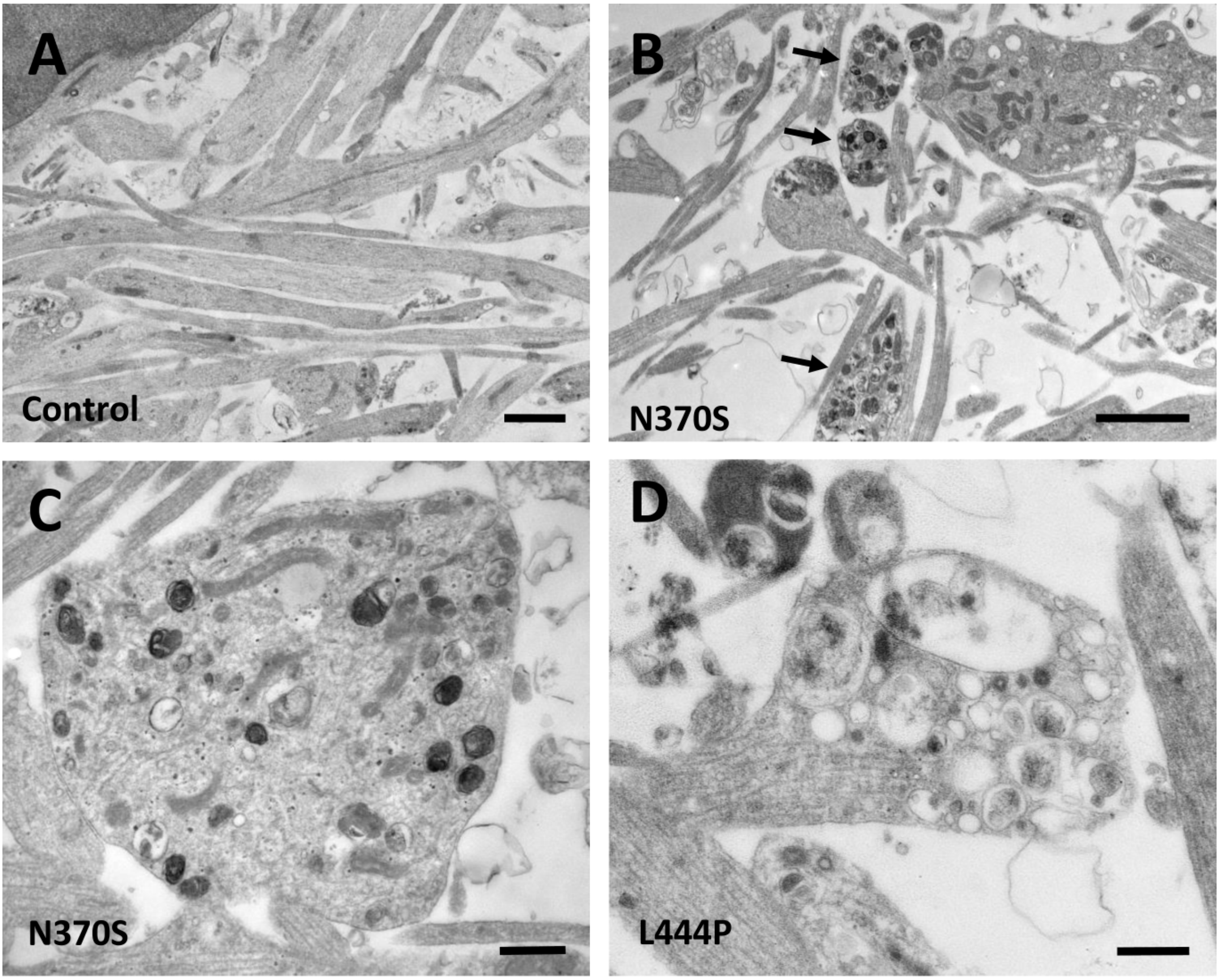

**Figure S7**

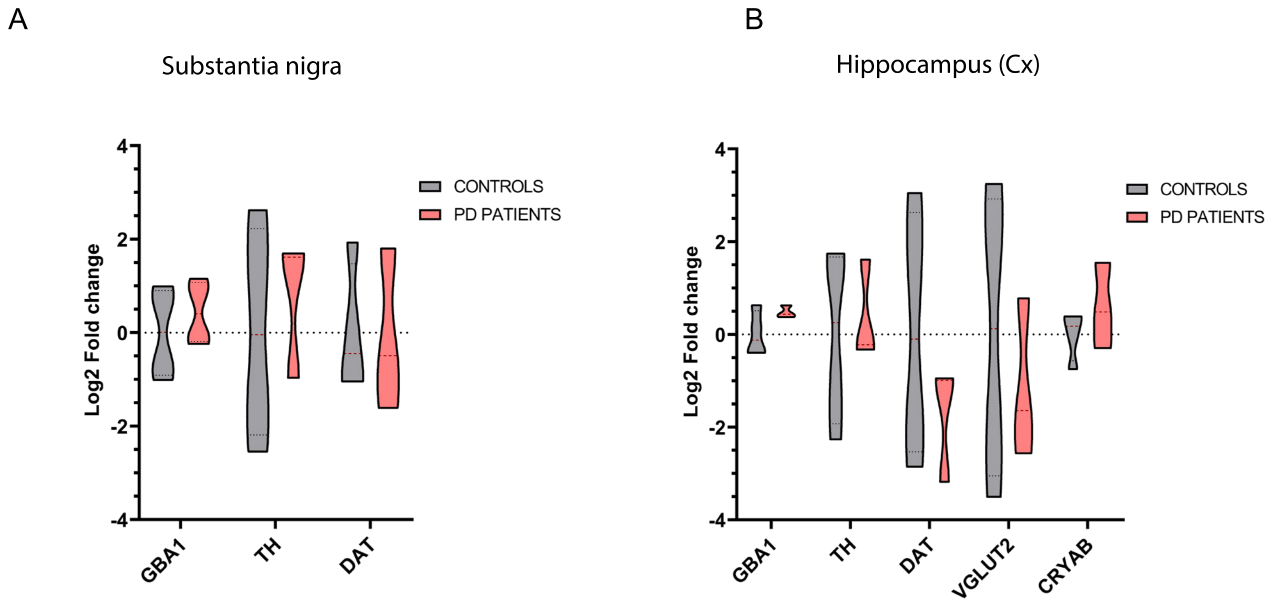
A
Substantia nigra
Log2 Fold change
4
2
0
-2
-4
CONTROLS
PD PATIENTS
GBA1
TH
DAT
B
Hippocampus (Cx)
Log2 Fold change
4
2
0
-2
-4
CONTROLS
PD PATIENTS
GBA1
TH
DAT
VGLUT2
CRYAB

| Table S1. Primers (5’-3’) used for the gene-expression analysis by RT-qPCR | | | |
|---|---|---|---|
| *CRYAB* | Forward | ATCCGCCGCCCCTTCTTTC | 181 bp |
| | Reverse | GGCGCATCTCTGAGAGTCCAGTG | |
| *FOXA2* | Forward | CCGGCCCATTATGAACTCCTCTTA | 144 bp |
| | Reverse | ATTTCTTCTCCCTTGCGTCTCTGC | |
| *GAPDH* | Forward | AACCATGAGAAGTATGACAACAGCC | 118 bp |
| | Reverse | TGAGTCCTTCCACGATACCAAAGT | |
| *GBA1* | Forward | CCACAGCATCATCACGAACCTCC | 154 bp |
| | Reverse | GGGCTGTTTGTAAAACGTGTCCTTG | |
| *GIRK2* | Forward | Izasa/Werfen (Cat#QT00010444) | 118 bp |
| | Reverse | | |
| *LMX1A* | Forward | CAGGGCTGAGTGTCCGTGTCG | 147 bp |
| | Reverse | CACTCCCACCACCGTTTGTCTGA | |
| *LMX1B* | Forward | TCCGTGAAGAGCGAGGATGAAGAT | 236 bp |
| | Reverse | TGGACCACGCGCACACTGAG | |
| *NURR1* | Forward | AGTTTAAAAGGCCGGAGAGGTCGT | 209 bp |
| | Reverse | AATTGCTGGATATGCTGGGTGTCA | |
| *PAX6* | Forward | CCCCAGCCAGACCTCCTCATACT | 185 bp |
| | Reverse | CGGGAACTTGAACTGGAACTGACAC | |
| *SLC17A6 (VGLUT2)* | Forward | CAAGGTTGGTATGCTATCTG | 118 bp |
| | Reverse | TGATCTTTCTCACTGTCGTAG | |
| *SLC18A2 (VMAT2)* | Forward | GCTATTCTCATGGATCACAACTGCC | 207 bp |
| | Reverse | CCAGGGATGGATGGTATGACTAAGAC | |
| *SLC6A3 (DAT)* | Forward | TCAGACCCCCCCACTACGGAG | 141 bp |
| | Reverse | TCTCTCGAAAGGACCCAGGCAG | |
| *TH* | Forward | CGGTGGAGTTCGGGCTGTGTAA | 217 bp |
| | Reverse | CTGACTTGTCCTTGGCGTCACT | |
| *TUBB3* | Forward | AAGTACGTGCCTCGAGCCATTCTG | 220 bp |
| | Reverse | GCAGGCAGTCGCAGTTTTCACAC | |
